# Investigating the Interaction Between EEG and fNIRS: A Multimodal Network Analysis of Brain Connectivity

**DOI:** 10.1101/2023.11.08.565955

**Authors:** Rosmary Blanco, Cemal Koba, Alessandro Crimi

**Affiliations:** Sano Centre for Computational Medicine, Czarnowiejska 36, C5, Krakow, 30-054, Poland; AGH University of Science and Technology of Krakow, Al. Mickiewicza 30, Krakow, 30-059, Poland

**Keywords:** EEG, fNIRS, Multimodal, Multilayer, Networks, Resting-state, Motor imagery

## Abstract

The brain is a complex system with functional and structural networks. Different neuroimaging methods have their strengths and limitations, depending on the signals they measure. Combining techniques like electroencephalography (EEG) and functional near-infrared spectroscopy (fNIRS) techniques has gained interest, but understanding how the information derived from these modalities is related remains an exciting open question. Successful integration of these modalities requires a sophisticated mathematical framework that goes beyond simple comparative analyses. The multilayer network model has emerged as a promising approach. This study is an extended version of the conference paper “Resting State Brain Connectivity Analysis from EEG and FNIRS Signals” [5]. In this study, we explored the brain network properties obtained from EEG and fNIRS data using graph analysis. Additionally, we adopted the multilayer network model to evaluate the benefits of combining multiple modalities compared to using a single modality. A small-world network structure was observed in the rest, right motor imagery, and left motor imagery tasks in both modalities. We found that EEG captures faster changes in neural activity, thus providing a more precise estimation of the timing of information transfer between brain regions in RS. fNIRS provides insights into the slower hemodynamic responses associated with longer-lasting and sustained neural processes in cognitive tasks. The multilayer approach outperformed unimodal analyses, offering a richer understanding of brain function. Complementarity between EEG and fNIRS was observed, particularly during tasks, as well as a certain level of redundancy and complementarity between the multimodal and the unimodal approach, which is dependent on the modality and on the specific brain state. Overall, the results highlight differences in how EEG and fNIRS capture brain network topology in RS and tasks and emphasize the value of integrating multiple modalities for a comprehensive view of brain connectivity and function.

## 1. Introduction

Contemporary neuroimaging methods depend on diverse source signals across various spatial and temporal scales. Neural activity is predominantly described by electrophysiological mechanisms and it is mediated by metabolic and hemodynamic processes. Unlike electrophysiological signals, metabolic and hemodynamic responses are much slower and reflect the indirect and secondary effects of neural activity. Noninvasive neuroimaging modalities are based on biophysical signals associated with either brain electrophysiology or hemodynamics/metabolism. The strengths and limitations of these modalities depend largely upon the spatiotemporal characteristics of the measured source signals in relation to neuronal activity, as well as many diverse sensing and imaging methods applied to individual modalities. The complementary features have motivated recent developments in integrating multiple neuroimaging modalities with a notable focus on the integration of EEG and fNIRS in recent years [47, 50, 26]. fNIRS relies on differential measurements of the backscattered light, which is sensitive to oxy-(HbO) and deoxyhemoglobin (HbR). EEG, on the other hand, captures the electrical brain activity derived from synchronous excitatory post-synaptic potentials. The former has high spatial resolution but is highly sensitive to scalp-related (extracerebral) hemoglobin oscillations. The latter allows the tracking of the cerebral dynamics with the temporal detail of the neuronal processes (1 ms) but suffers from volume conduction. Their integration holds the promise of achieving spatial resolutions on the order of millimeters and temporal resolutions in the range of milliseconds, representing an exclusive opportunity to explore the dynamics of brain activity. In this context, network neuroscience offers a valuable approach to model and investigate brain dynamics, as well as to explore the potential of multimodal approaches in inferring brain functions. Large-scale functional brain connectivity can be modeled as a network or graph. When different modalities are examined, it is common to perform global comparisons [42, 58], which allows direct comparison of the properties of the different modalities. Regional and network-level relationships have been explored using electrophysiological and fMRI recordings, suggesting that the relationship between the two modalities may be affected by distinct cytoarchitecture and laminar structure of brain regions [46]. Moreover, the relationship between hemodynamic and electromagnetic networks has been predominantly focused within different frequency bands, suggesting that hemodynamic connectivity cannot be explained by electrical activity in a single frequency band but rather arises from the combination of multiple frequency bands [45]. However, how the electromagnetic and hemodynamic networks relate to one another remains an exciting open question. Given the increasing interest in multimodal imaging approaches, there is a need for a more sophisticated mathematical framework exceeding merely comparative analyses between modalities. Such a framework should discern the similarities or dissimilarities between the functional EEG and fNIRS networks. Within this context, the graph-based network analysis approach is a powerful tool for characterizing topological properties of the brain networks. In addition, the multilayer network approach has emerged as an innovative mathematical framework that facilitates the integration of different biophysical information, displaying promising outcomes in the field of neuroscience [59, 13, 29]. The primary contribution of this study is to investigate the correspondence between fNIRS-based functional connectivity and EEG functional connectivity, emphasizing the shared patterns and distinctions in the network structures of the two modalities through graph analysis. Additionally, we shed light on how the interplay between electrophysiological and hemodynamic activities varies between the resting state (RS) and task-related conditions. Lastly, we provide a more holistic perspective on brain functionality by using a multilayer approach to investigate whether the integration of complementary information from various sources provides a clearer picture of brain dynamics than the unimodal approach for inferring brain functions.

## 2. Materials and Methods

Synchronous EEG and fNIRS recordings during the resting state and motor imagery task from 18 healthy subjects (28.5 ± 3.7 years) were obtained from an open dataset [49]. EEG data were recorded with 32 electrodes placed according to the international 10-5 system (AFp1, AFp2, AFF1h, AFF2h, AFF5h, AFF6h, F3, F4, F7, F8, FCC3h, FCC4h, FCC5h, FCC6h, T7, T8, Cz, CCP3h, CCP4h, CCP5h, CCP6h, Pz, P3, P4, P7, P8, PPO1h, PPO2h, POO1, POO2 and Fz for ground electrode). Those are referenced to the linked mastoid at a 1000 Hz sampling rate (down-sampled to 200 Hz). fNIRS data were collected by 36 channels (14 sources and 16 detectors with an interoptode distance of 30 mm), following the standardized 10-20 EEG system, at a 12.5 Hz sampling rate (down-sampled to 10 Hz) as depicted in Figure 1. Two wavelengths at 760 nm and 850 nm were used to measure the changes in oxygenation levels. all the steps of the pipeline, starting from EEG and fNIRS data acquisition to the construction of the brain network, along with an overview of the methods used are summarized in Figure 2.

**Figure 1:**
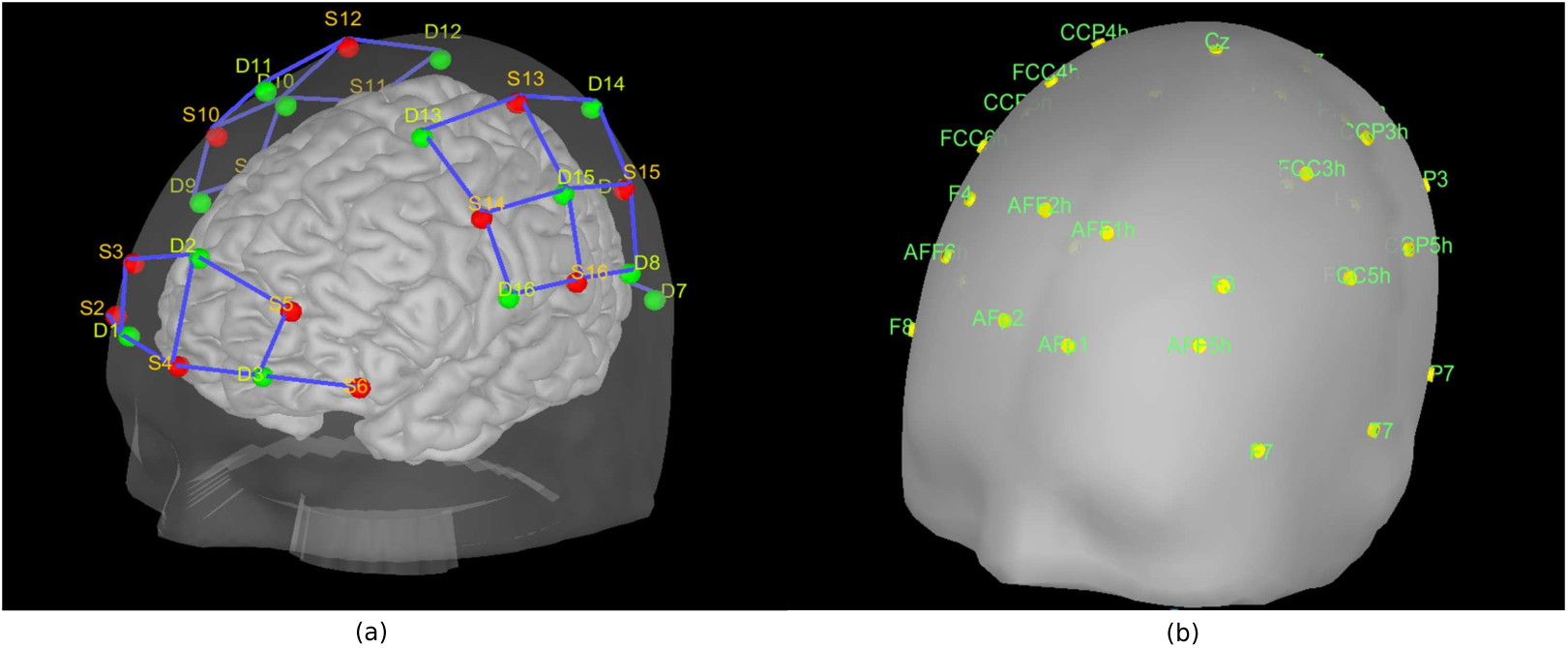
(a): NIRS optode locations. The red dots are the sources and the green dots are the detectors. (b): EEG electrode locations.

**Figure 2:**
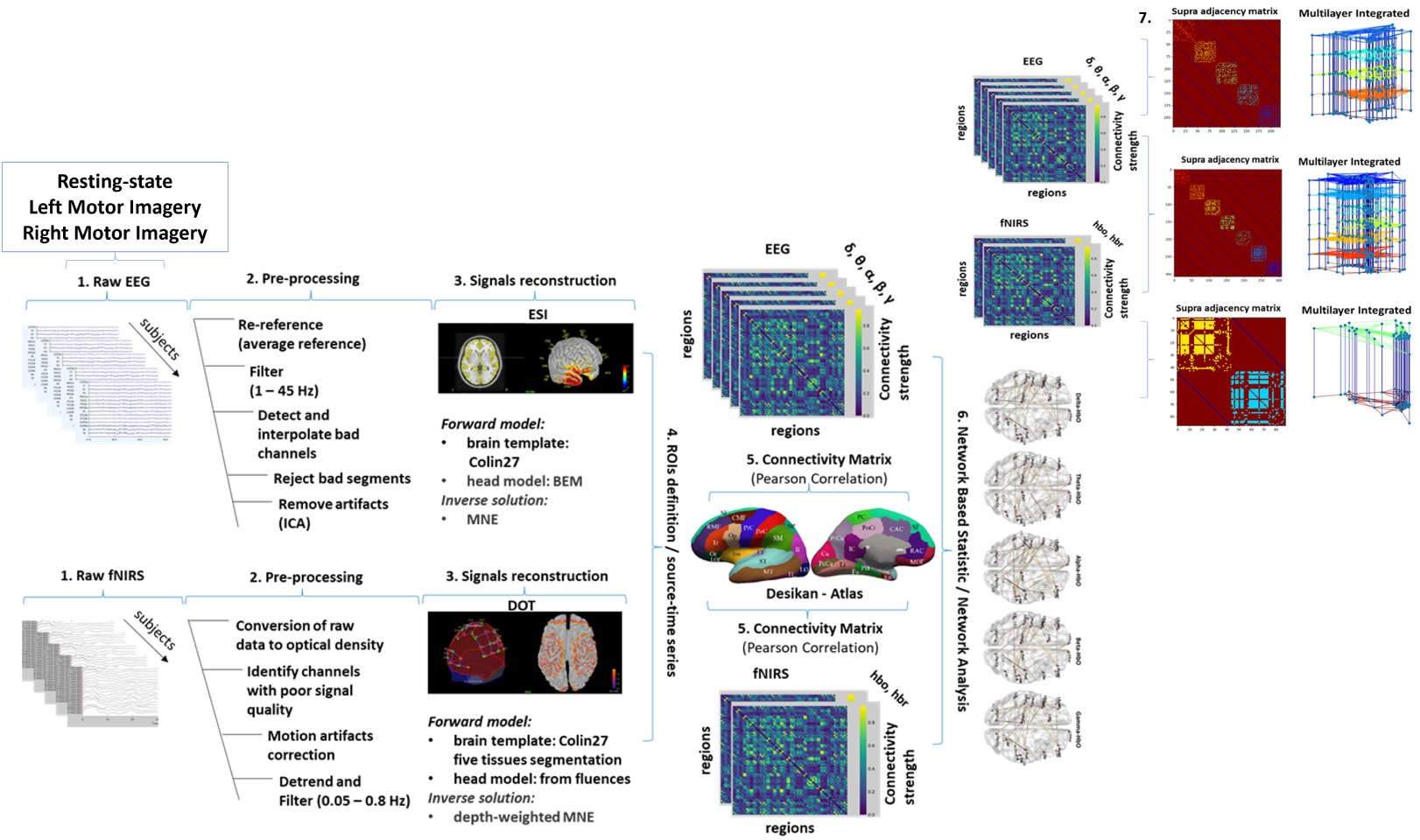
Method workflow. 1. The EEG and fNIRS data were collected during the resting state (RS), left motor imagery (LMI), and right motor imagery (RMI). 2. The pre-processing steps for cleaning data. 3. The methods for the reconstruction of the signals in the source space. The Electrical Source Imagine (ESI) technique to estimate the cortical sources (neuronal activity) by segmenting the MRI and building a realistic head model (forward model). This model is used to solve the inverse problem for the estimate of the amplitude of the current dipoles captured by the sensors on the scalp (differences in electrical potentials recorded by the electrodes). The Diffuse Optical Tomography (DOT) technique to model light propagation within the head and estimate the optical properties of each tissue (forward model) by fluences of light for each optode (source and detector). This model maps the absorption changes along the cortical region (source space) to the scalp sensors (measured as optical density changes). By solving the inverse problem the cortical changes of hemodynamic activity within the brain are estimated. 4. The EEG and fNIRS source-time series were mapped in the same 3D space using an atlas-based approach (Desikan-Killiany), with the definition of Region of Interest (ROIs) at which both signals were estimated. 5. Functional connectivity (Pearson’s correlation for fNIRS, PLV for EEG) estimates the statistical coupling between each ROI pair of the reconstructed time series. 6. The topology of brain networks captured by the two techniques was compared through graph theoretical approaches. The similarities and differences in connectivity strength and patterns between EEG and fNIRS networks were identified by means of the cluster-based thresholding approach. 7. The multilayer approach is applied to investigate whether the integration of the information outperforms the unimodal approach in characterizing brain network topology and assess if the combined information provides complementary or redundant information compared to a single modality, employing graph analysis.

### 2.1. EEG and fNIRS data pre-processing

Compared to our previous study [5], we have introduced further preprocessing steps aimed at enhancing data quality, primarily for fNIRS signals. Following the implementation of this new preprocessing pipeline (details are below), only 18 subjects out of the initial 29 have been retained as they exhibited high-quality signals in both EEG and fNIRS and were thus included in the subsequent analyses. A standardized pre-processing pipeline was applied to remove artifacts from EEG and fNIRS recordings using the MNE toolbox [22]. EEG data was first re-referenced using a common average reference and filtered (second-order zero-phase Butterworth type FIR low-pass filter) with a cut-off of 1 Hz. The infomax algorithm was employed for performing independent component analysis (ICA) in a semi-automatic manner to identify and eliminate artifacts originating from biological sources such as muscle activity, blinks, and eye movements, as well as non-biological sources such as line noise and other environmental or instrumental interferences [39]. This involved visually inspecting various diagnostic measures of each independent component, including topography, epochs image, power spectrum, and epoch variance. Then the cleaned data were filtered (second-order zero-phase Butterworth type FIR highpass filter) with a cut-off of 45 Hz. The implemented pre-processing pipeline for fNIRS data included the conversion of raw data to optical density (OD) and the assessment of data quality through the computation of the scalp-coupled index (SCI). We then identified the channels displaying an SCI *<* 0.7, which were labeled as ‘bad channels’. Subjects exhibiting more than 50% of channels marked as bad were excluded. The OD signals were filtered by a finite impulse response (FIR) bandpass filter with cutoff frequencies of 0.02-0.08 Hz. Bad time segments containing excessive head movements were identified and rejected using the global variance in temporal derivative (GVTD) metric, inspired by the concept of Derivative of Variance of Root Mean Square (DVARS) in fMRI [48]. Time points with GVTD values larger than three times the mean GVTD value across all time points were flagged, and 15-second time windows centered around these points were considered bad time segments. These segments were automatically annotated and removed from the recordings. In order to address the issue of systemic physiological effects on the fNIRS signals, we decomposed the OD signals into multiple components by principal component analysis (PCA) and evaluated the spatial uniformity of each principal component by the metric of coefficient of spatial uniformity ([31]). Given that high spatial uniformity is indicative of superficial skin responses ([62]), the principal component with the highest spatial uniformity value was identified and subsequently removed from the reconstructed signals.

### 2.2. Signal reconstruction

Source EEG and fNIRS data reconstruction in source space were performed using Brainstorm software [55] and custom Matlab scripts [2]. For EEG a multiple-layer head model (Boundary Element Method-BEM) and an MRI template available in BrainStorm (Colin27) were used to build a realistic head model (forward model) by the OpenMEEG tool [21], which takes the different geometry and conductivity characteristics of the head tissues into account. The dipoles corresponding to potential brain sources were mapped to the cortical surface. The surface map was parcellated with a high-resolution mesh (15000 vertices). Finally, a lead field matrix, expressing the scalp potentials corresponding to each single-dipole source configuration, was generated based on the volume conduction model [34]. For the fNIRS, a five-tissue segmentation of the Colin27 brain template was used for the sensitivity matrix (forward model) computation. The fluences of light for each optode were estimated using the Monte Carlo simulations with a number of photons equal to 10^8^ and projecting the sensitivity values within each vertex of the cortex mesh. The sensitivity values in each voxel of each channel were computed by multiplication of fluence rate distributions using the adjoint method according to the Rytov approximation [53]. Then the NIRS forward model was computed from fluences by projecting the sensitivity values within each vertex of the cortex mesh using the Voronoi-based method, a volume-to-surface interpolation method that preserves sulci-gyral morphology [27]. Given each forward model for EEG and fNIRS, the brain activity was estimated by the mathematical formulation of the inverse solution using the Minimum Norm Estimate (MNE) method. The minimum norm estimator is defined as:

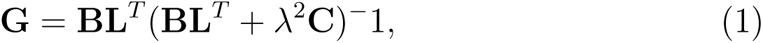

where G represents the reconstructed signal along the cortical surface, L is the non-zero element matrix inversely proportional to the norm of the lead field vectors, B is the diagonal matrix derived from L, capturing the weight of each source, C is the noise covariance matrix describing the statistical properties of noise, and *λ* is the regularization parameter that controls the trade-off between data fidelity and noise suppression, which is crucial for obtaining meaningful and accurate reconstructions. For EEG data, we employed the standardized Low-Resolution brain Electromagnetic Tomography (sLORETA) to apply the MNE method. In the case of fNIRS data, we utilized the depth-weighted minimum norm estimate (depth-weighted MNE) method. This choice was necessary because the standard MNE method tends to bias the inverse solution toward more superficial sources due to its sensitivity to near-surface signals. The depth-weighted MNE method addresses this bias by accounting for the varying light sensitivity at different depths within the brain tissue.

### 2.3. Unimodal network construction

In order to be able directly compare the EEG and fNIRS source-time series, we mapped them to the same 3D space using an atlas-based approach (Desikan-Killiany) [17] identifying 44 Regions of Interest (ROIs) that cover both modalities. The EEG ROI time series were then decomposed into the typical oscillatory activity by band-pass filtering: *δ*(1 *−* 4*Hz*)*, θ*(4 *−* 7*Hz*)*, α*(8 *−* 15*Hz*)*, β*(15 *−* 25*Hz*), and *γ*(25 *−* 45*Hz*), while fNIRS source time series were converted into oxygenated hemoglobin (HbO) and deoxygenated hemoglobin (HbR) by applying the modified Beer-Lambert transformation [10]. The Functional Connectivity (FC) matrices for the 44 ROIs within EEG (for each frequency band) were derived through the utilization of Phase Locked Value (PLV), whereas for fNIRS (HbO and HbR) Pearson’s correlation coefficient was employed to compute the FC matrices. This process generated a total of 7 connectivity matrices for each subject (5 for each EEG frequency band and 2 for hemodynamic activity of fNIRS) and for each condition (RS, RMI, and LMI), resulting in a total of 21 FCs.

To facilitate the comparison and ensure that the thresholding process is more consistent across modalities, we first normalized the edge weights in both EEG and fNIRS FCs, and then, given our objective of investigating distinct topological features in brain networks obtained from EEG and fNIRS separately, we chose a thresholding method based on the percolation theory. This thresholding method has the advantage of preserving the topological properties of the original weighted networks by taking into account the whole network topology during the binarization process. We used the maximum threshold that maintains the topological integrity of the original network, which in this case is defined as the size of the largest connected component in the network [18].

### 2.4. Connectivity analysis of unimodal networks

#### 2.4.1. Small-wordness

We compared the topological characteristics of the group-level functional brain network derived from EEG and fNIRS for the three conditions (RS, RMI, and LMI) through *graph/network analysis* using the brain connectivity toolbox (BCT) [44, 61]. To characterize the global topological organization of the whole-brain network, we chose Small-World propensity (*ϕ*), Global Efficiency (GE), Global Clustering Coefficient (GCC), and characteristic Path Length (PL) as features of interest.

The small-world structure of a network characterizes its efficiency and cost-effectiveness. The mathematical formulation of the small-world index (*σ*) is represented as 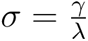, which indicates small-world characteristics when *σ >* 1. However, since *σ* is defined as a ratio, when *γ ≫* 1 and *λ >* 1, networks will consistently show *σ >* 1. To overcome this limitation, we decided to adopt the metric of the small-world propensity. Specifically, the small-world propensity, denoted as *ϕ*, quantifies how much a network’s clustering coefficient (*C_brain_*) and characteristic path length (*L_brain_*) deviate from those of lattice (*C_lattice_*, *L_lattice_*) and random (*C_random_*, *L_random_*) networks constructed with the same number of nodes and the same degree distribution:

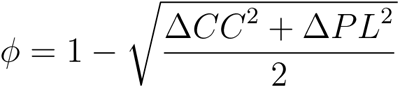

Where:

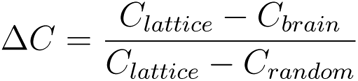

and

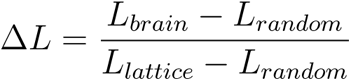

Networks are considered small-world if they have a small-world propensity 0.4 *< ϕ ≤* 1. The concept of small-world propensity addresses several limitations found in previous scalar definitions of small-worldness. Firstly, it can incorporate weighted evaluations of the clustering coefficient and path length. Secondly, it is independent of network density, allowing for comparisons of small-worldness between networks with significantly different densities. Lastly, this metric is informed by spatially-constrained null models [3].

#### 2.4.2. Global graph metrics

Among the global graph metrics, as a measure of functional integration, the GE is defined as the efficiency of information exchange in a parallel system, in which all nodes are able to exchange information via the shortest paths simultaneously. Its inverse, the path length, is the average shortest path length between all pairs of nodes in the network. As a measure of functional segregation, the CC is the fraction of triangles around an individual node reflecting, on average, the prevalence of clustered connectivity around individual nodes.

The graph measures were normalized by generating 100 randomized networks (null models of a given network) while preserving the same number of nodes, and degree distribution. Then, for each measure, the ratio was calculated as *real metric* over the *matched random metrics*. For the comparison of network topology features, we utilized the Wilcoxon test for paired data to assess the differences between EEG frequency bands with their corresponding fNIRS metrics, specifically (*δ, θ, α, β, γ* - HbO) and (*δ, theta, α, β, γ* - HbR). Significance was established at a threshold of p *<* 0.05. To account for multiple comparisons, we applied the False Discovery Rate (FDR) correction.

#### 2.4.3. Edge-wise analysis

To assess the similarities and differences in connectivity strength and patterns between EEG and fNIRS networks, we employed an *edge-wise analysis*. Significant connections were identified by means of the cluster-based thresholding approach using the brain connectivity toolbox (BCT) [44, 61]. For each edge of the 44 x 44 connectivity matrix, a two-sample paired t-test was performed independently between each modality (5 frequency bands and 2 hemodynamic responses), and cluster-forming thresholds were applied to form a set of suprathreshold edges. The threshold was chosen based on Hedge’s g-statistic effect size (ES) computed between each node of the two matrices, pairing each EEG frequency band FC with each fNIRS FC. The t-statistic corresponding to the Hedge’s g score closest to 0.5 (medium ES) was chosen as the critical value (t-stat = 2.5, corresponding to p-values = 0.01). Finally, an FWER-corrected p-value was ascribed to each component through permutation testing (5,000 permutations). Edges that displayed FWER-corrected p-values below the significance threshold of 0.05 were considered positive results.

We employed both a right-tailed t-test and a left-tailed t-test because of the absence of a priori assumptions regarding the direction of differences between the two EEG and fNIRS networks. The right-tailed test investigates whether the EEG network is greater than the fNIRS network, while the lefttailed test explores the possibility of the opposite relationship.

Additionally, to assess whether the resulting significant networks exhibited similarities across different frequency bands, we computed the Pearson correlation coefficient to determine the similarity between each pair of resulting matrices. For this purpose, a threshold of r = 0.5 was chosen arbitrarily as the criterion for defining such similarity, without optimizing this value. If two matrix pairs displayed an r-value *<* 0.5, it indicated that the relationship between the hemodynamic and electrophysiological signals varied depending on the specific frequency band.

### 2.5. Multimodal network construction

In order to assess the potential advantages of integrating information from different modalities over using a single modality, we opted for a multilayer network approach composed using custom-made scripts that integrate the Python libraries multiNetX [51] and NetworkX [24]. This involved constructing a multiplex network of integrated EEG and fNIRS networks. The multiplex is a multilayer network employed to depict various interactions among the identical set of nodes. Each layer is distinguished by a distinct mode of interaction, and the interlayer connections exclusively link the same node across all layers. This mathematical framework proves valuable for encoding information from brain networks generated using diverse edge weights as all layers are built using the same atlas. To mitigate potential bias due to the differences in link density and average connectivity across layers, we adopted a data-driven approach for the thresholding of the original EEG and fNIRS FCs, which were utilized in the construction of the multiplex networks. Specifically, we generated binary networks in the shape of Minimum Spanning Trees (MST) by applying Kruskal’s algorithm to the functional connectivity matrices associated with each EEG frequency band, as well as the fNIRS HbO and HbR data, as depicted in [6]. The MST is supposed to reduce the influence of spurious connections within the brain by considering strong connections that don’t form cycles or loops to enhance the specificity of network topological characteristics. This configuration serves as the backbone of the network and is unaffected by prevalent methodological issues including potential impacts of connection strength or link density on the inferred topological attributes of networks [52, 56].

We integrated the MSTs to obtain an interconnected multiplex network (Full) for every subject. Each participant’s multiplex thus consisted of L = 7 layers: two for fNIRS (HbO and HbR) and five for EEG (*δ, θ, α, β, γ*), with each layer containing the same set of N = 44 nodes, and M = N – 1 = 43 intralayer links. The weights of the interlayer connections were set to 1, identical to the intralayer connections, because there is no established method for determining biologically meaningful weighted interlayer links between different modalities [6]. The resulting multilayer network was an LxN by LxN supra-adjacency matrix (Figure 3) with diagonal blocks encoding intralayer connectivity for each modality and off-diagonal blocks encoding interlayer connectivity. This multilayer was built for the three conditions (RS, LMI, and RMI), resulting in three multilayer networks. We then applied the same procedure to build the multiplex network exclusively for EEG (L = 5 layers) by integrating the 5 FCs (*δ, θ, α, β, γ*), and for fNIRS by integrating the 2 HbO and HbR FCs (L = 2 layers). These multilayers were built for the three conditions, resulting in three multilayer networks for EEG and three multilayer networks for fNIRS.

**Figure 3:**
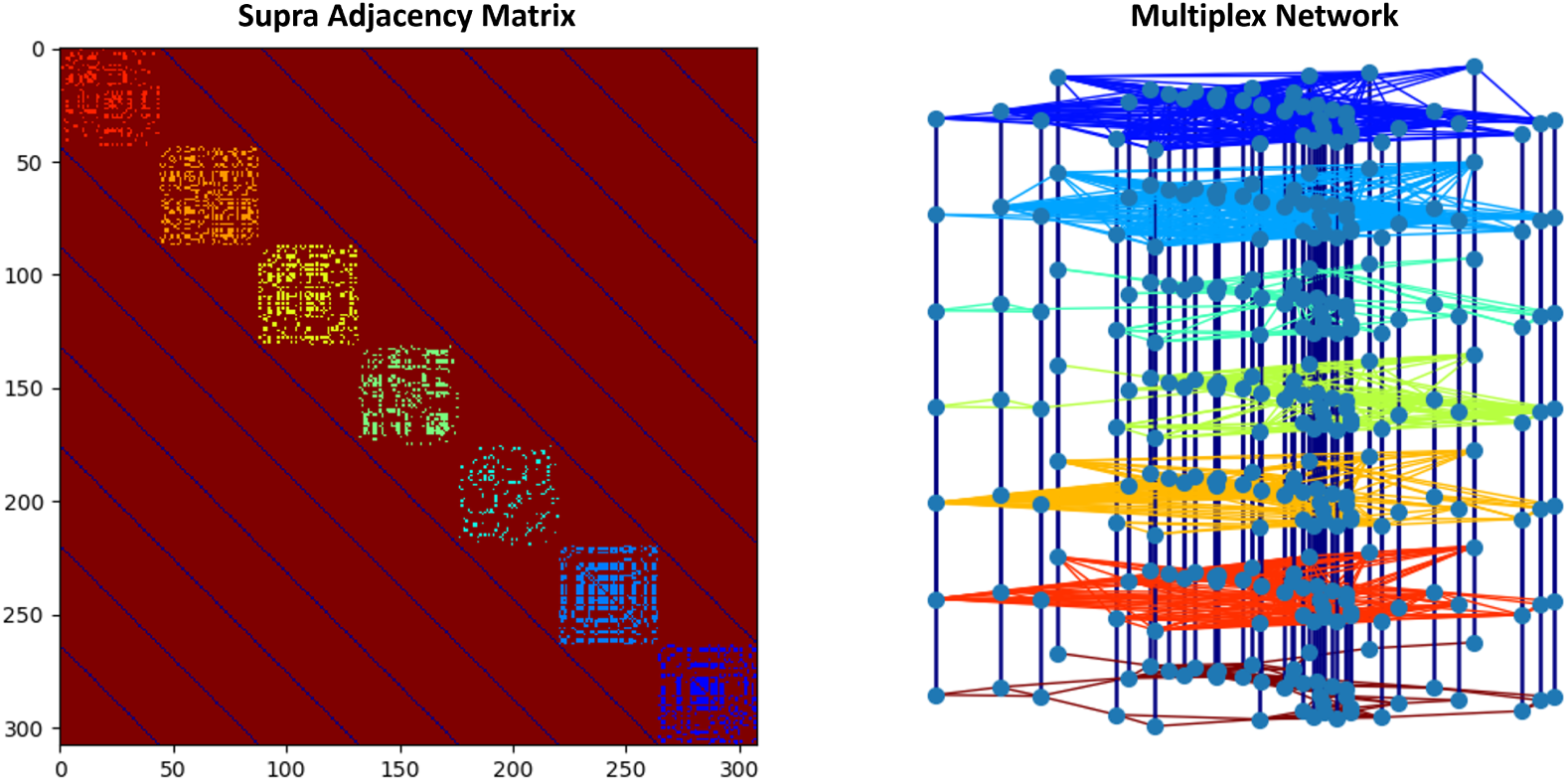
Supra-adjecency matrix

### 2.6. Connectivity analysis of multimodal networks

To investigate whether the integration of the information derived from different sources outperforms the unimodal approach in characterizing the topology of the whole-brain network, we compared the full-multilayers with the EEG- and fNIRS-multilayers by means of graph analysis. We chose the global multilayer graph metrics of Modularity (M), Global Efficiency (GE), and Path Length (PL) as measures of the effectiveness of the communication, as those are the most commonly reported in the literature. M in graph theory refers to a measure that quantifies the division of a network into modules or communities, where nodes within the same module are more densely connected to each other than nodes in different modules. It provides insights into the functional subgroups present in the network. High M values indicate a strong tendency for nodes to cluster together, while low values suggest a more random or uniform distribution of connections. All the global graph metrics were computed for each subject by aggregating these measures across layers to obtain one value per subject. A paired t-test for was used to assess the differences between full-multilayers and EEG-multilayers and between full-multilayers and fNIRS-multilayers network features. Significance was established at a threshold of p *<* 0.05.

Additionally, we investigated whether the integrated information in the full-multilayer provides complementary or redundant information compared to a single modality (EEG-multilayer or fNIRS-multilayer). To achieve this, we calculated the nodal metric of Eigenvector Centrality (EC) to identify key nodes in the network, thus characterizing brain patterns. For each subject, the EC values were computed for each node in each layer separately and subsequently aggregated across layers to obtain one value per node, as described in [15]. We evaluated the (dis)similarity of brain activation patterns between the full-multilayers and each EEG- and fNIRS-multilayers using the Dice coefficient of similarity, a statistic used to gauge the similarity of two samples, often used in image segmentation. For the calculation of this coefficient, we averaged nodal EC across subjects to obtain one value for each node per multilayer. This allowed us to assess the extent to which the multimodal approach aligns with the unimodal approaches in evaluating central brain regions during both resting state and task conditions. The high similarity suggests that, in a particular condition, information integration has a similar effect on capturing network structure compared to a single modality, indicating redundancy. Conversely, low similarity suggests that the integration of different modalities leads to differences in network structure, emphasizing the presence of complementary information.

Then, we systematically computed differences in each task condition relative to the RS as a baseline, enabling us to capture variations in brain activity during LMI and RMI for the Full-Multilayer, EEG-Multilayer, and fNIRS-Multilayer, separately. Subsequently, we extracted the cortical regions linked to the MI task according to the literature. These regions encompass specialized areas involved in motor planning and execution processes during MI, including the motor cortex (PreC), supplementary motor cortex (PaC), and somatosensory cortex (PoC). We also considered regions associated with MI, such as the temporo-occipital association areas (Cu, Fu, and MT), which are engaged in various cognitive functions, including attention, memory, and mental simulation of actions; the frontal and prefrontal associative cortex premotor cortex (SF) and dorsolateral prefrontal cortex (RoMF and CMF), involved in higher-order cognitive processes, including motor planning, memory, and decision-making; the associative parietal areas (IP, SP, and SM), which contributes to various functions related to spatial awareness and sensorimotor integration.

## 3. Results

### 3.1. Unimodal networks

#### 3.1.1. Small-worldness

The primary aim of the study was to investigate the relationship between fNIRS-based functional connectivity and EEG functional connectivity, with a focus on identifying commonalities and differences in their network structures. To achieve this we compared the topological characteristics of the group-level functional connectivity derived from EEG and fNIRS for the three conditions: RS, RMI, and LMI. For the assessment of the global topological organization of the whole-brain network, the small-world propensity (*ϕ*), Global Efficiency (GE), global Clustering Coefficient (GCC), and characteristic Path Length (PL) metrics were chosen. Topology analyses showed that all the EEG frequency bands (*δ, θ, α, β, γ*) and fNIRS (HbO and HbR) have 0.4 *< ϕ ≤* 1, which implies prominent small-world properties in both modalities and for each condition (Figure 4a). A real network would be considered to have the small world characteristics if CC*_real_*/CC*_rand_ >* 1 and PL*_real_*/PL*_rand_ ≈* 1. It means that compared to random networks, a true human brain network has a larger GCC and an approximately identical PL between any pair of nodes in the network. This was demonstrated for all EEG frequency bands by showing significantly higher GCC values than HbO (Figure 4b). It means EEG networks have better clustering ability and smallworldness than fNIRS. In particular, the *α* band exhibited higher clustering in the RS condition, where this brainwave predominates. Conversely, during RMI and LMI *β* and *γ* bands showed higher GCC values, emphasizing the significance of these brainwaves in characterizing MI tasks.

**Figure 4:**
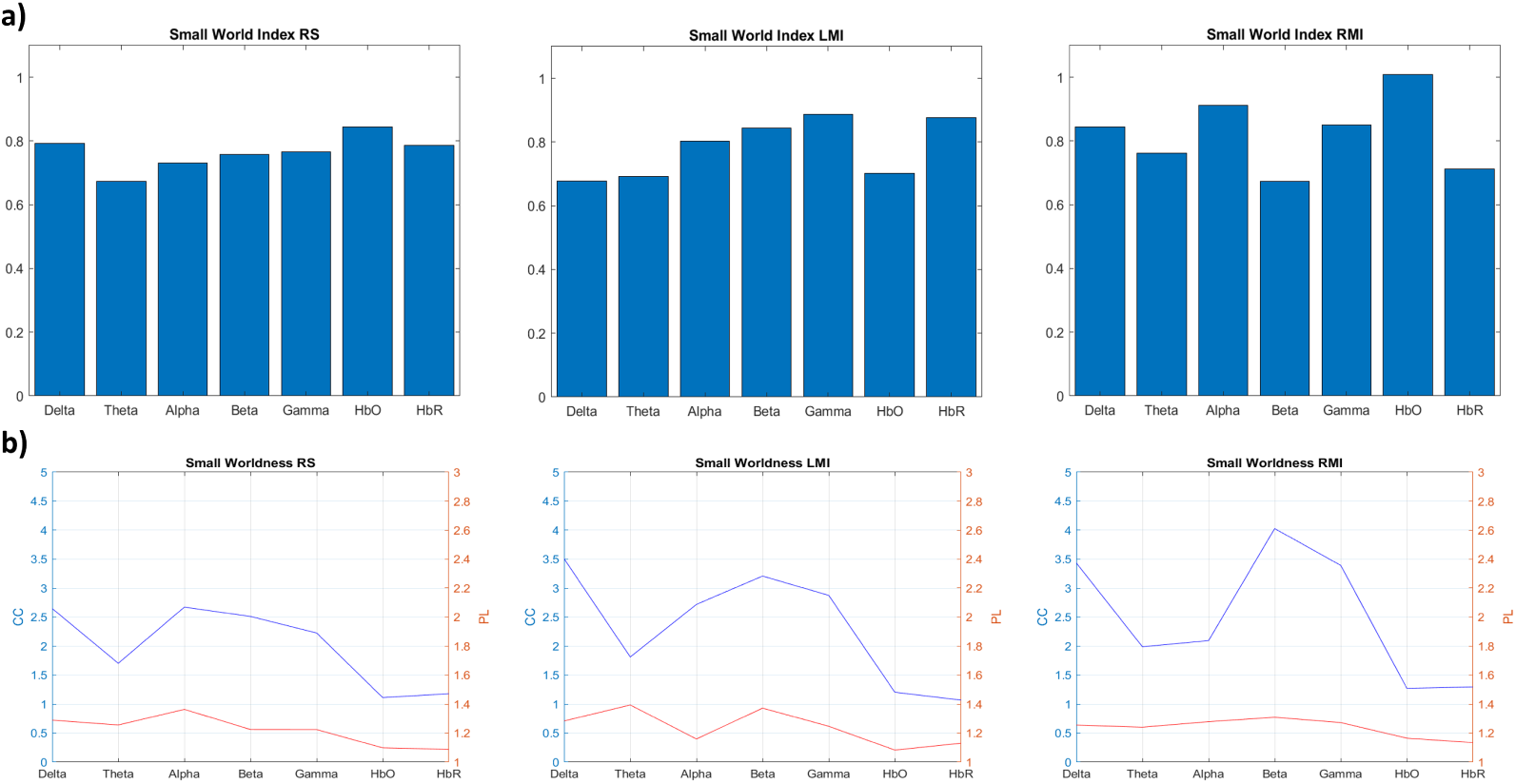
a) Barplot of Small-World propensity (*ϕ*) across EEG (Delta, Theta, Alpha, Beta, Gamma) and fNIRS networks, b) plot of the normalized Characteristic Path Length (PL) and the normalized Global Clustering Coefficient (GCC) for EEG (Delta, Theta, Alpha, Beta, Gamma) and fNIRS (HbO and HbR).

#### 3.1.2. Global graph metrics

Our observations revealed notable variations in GE, PL, and GCC values across different frequency bands and hemoglobin types (Figure 5, Table 1). We observed significantly higher GE values in the *θ* and *β* bands compared to HbO, with no statistical differences in the LMI and RMI conditions. When comparing to HbR, higher GE values were evident in the *θ* band during RS, and in both the *δ* and *θ* bands during the RMI condition. No significant differences were observed in the LMI condition. During RS, EEG shows higher information exchange capacity in specific frequency bands compared to fNIRS, while in task conditions, the efficiency of distributed information processing in the brain network between the two modalities is more similar. For the PL measure, we observed significantly lower values in RS for *δ, θ, β*, and *γ* bands compared to HbO, and *θ, β* and *γ* bands compared to HbR. In RMI, *δ, θ, α*, and *γ* bands exhibited lower values compared to HbO, and all the frequency bands had lower values compared to HbR. For LMI, HbO and HbR showed differences in the *δ, α*, and *γ* bands for fNIRS *>* EEG. The variations observed in these frequency bands further emphasize the complex interplay of modalities and conditions in shaping the brain’s functional connectivity and network efficiency. Statistically higher values for GCC were exhibited in *δ, α, β*, and *γ* bands compared to HbO and all the frequency bands vs. HbR during the RS. Brain regions within these frequency bands are more likely to form tightly connected clusters, suggesting a higher degree of local connectivity or modularity during rest. During the task, for LMI, higher GCC values were shown in *δ, α, β*, and *γ* compared to both HbO and HbR, while for RMI, in *δ, β*, and *γ* bands compared to both HbO and HbR. This finding implies that these frequency bands play a significant role in the local connectivity patterns within the brain network during cognitive tasks.

**Figure 5:**
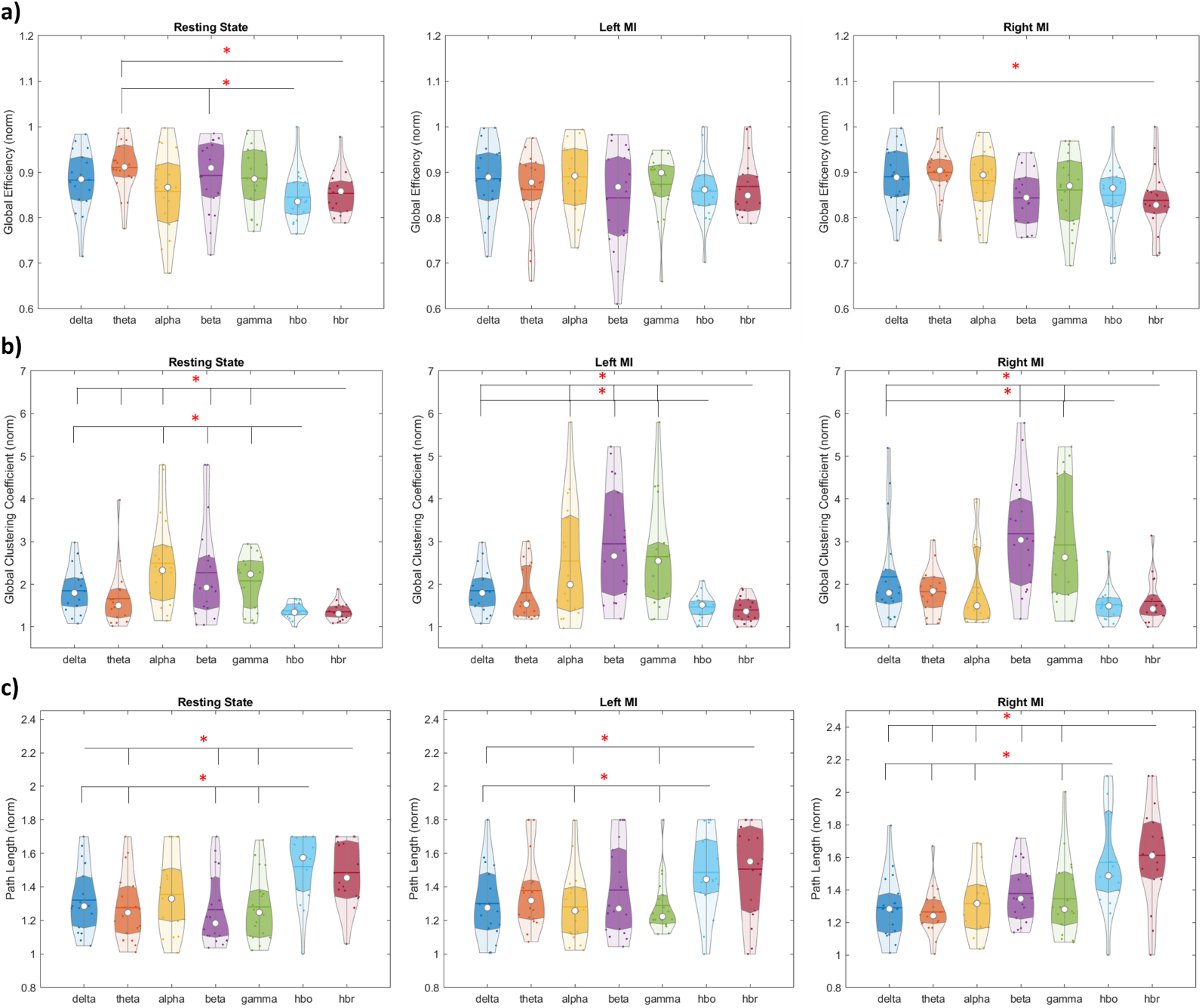
Violinplot of (a) Global Efficiency (GE), (b) Characteristic Path Length (PL) and (c) Global Clustering Coefficient (GCC) for EEG (Delta, Theta, Alpha, Beta, Gamma) and fNIRS (HbO and HbR). The red asterisk denotes the significant difference between EEG and fNIRS (p-value *<* 0.05, FDR corrected).

**Table 1:** Z scores of the Wilcoxon tests for Global Efficiency (GE), Characteristic Path Length (PL) and Global Clustering Coefficient (GCC) for EEG (Delta, Theta, Alpha, Beta, Gamma) and fNIRS (HbO and HbR)

|  | Alpha-HbO | Delta-HbO | Theta-HbO | Beta-HbO | Gamma-HbO | Alpha-HbR | Delta-HbR | Theta-HbR | Beta-HbR | Gamma-HbR |
| --- | --- | --- | --- | --- | --- | --- | --- | --- | --- | --- |
| **RMI** |  |  |  |  |  |  |  |  |  |  |
| GE | 1.546 | 1.59 | 1.894 | -0.414 | 0.936 | 1.894 | 2.286 | 2.548 | 0.24 | 0.936 |
| GCC | 1.459 | 2.112 | 2.156 | 3.68 | 3.593 | 1.111 | 1.72 | 1.59 | 3.375 | 3.375 |
| PL | -2.722 | -2.722 | -2.896 | -1.851 | -2.243 | -2.853 | -2.983 | -3.332 | -2.243 | -2.373 |
| **LMI** |  |  |  |  |  |  |  |  |  |  |
| GE | 1.372 | 1.154 | 0.675 | -0.457 | 1.067 | 1.111 | 0.414 | -0.283 | -1.023 | 0.501 |
| GCC | 2.504 | 1.982 | 1.59 | 3.332 | 3.114 | 2.896 | 2.243 | 1.72 | 3.68 | 3.506 |
| PL | -2.069 | -2.199 | -1.328 | -0.849 | -2.025 | -2.112 | -2.678 | -0.762 | -0.588 | -2.243 |
| **RS** |  |  |  |  |  |  |  |  |  |  |
| GE | 0.762 | 1.677 | 2.548 | 1.982 | 1.677 | 0.152 | 1.415 | 2.548 | 1.677 | 1.546 |
| GCC | 3.245 | 3.419 | 1.241 | 2.765 | 3.114 | 3.506 | 2.853 | 2.112 | 3.157 | 3.245 |
| PL | -1.546 | -2.548 | -2.765 | -2.504 | -2.765 | -1.154 | -1.807 | -2.33 | -2.765 | -2.417 |

#### 3.1.3. Edge-wise differences

The edge-wise analysis was applied via the cluster-based thresholding approach to assess the similarities and differences in connectivity strength and patterns between EEG and fNIRS networks. This analytical method identified subnetworks characterized by increased functional connectivity in HbO and HbR compared to EEG, as determined by a left-tailed t-test, across all frequency bands and conditions (RS, LMI, and RMI) at a predefined thresh-old of 2.5. Specifically, during the RS condition, we identified a large subnetwork with increased connectivity in *δ*-HbO, *α*-HbO, and *β*-HbO (Figure 6), and in *δ*-HbR, *θ*-HbR, *α*-HbR, *β*-HbR, and *γ*-HbR (Figure 7). This subnetwork exhibited preferentially interhemispheric and homotopic connections between various cortical regions, including the left precuneus (Cu), bilateral inferior parietal (IP), bilateral supplementary motor (PaC), post-postcentral gyrus (PoC), right middle temporal gyrus (MT), left medial frontal cortex (mOF), and bilateral pars opercularis of inferior frontal gyrus (Op). While, for *θ*-HbO and *γ*-HbO, two distinct subnetworks were identified. One subnetwork resembled the aforementioned large subnetwork, while the other comprised the right superior frontal cortex (SF), superior parietal (SP), and superior temporal regions (ST).

**Figure 6:**
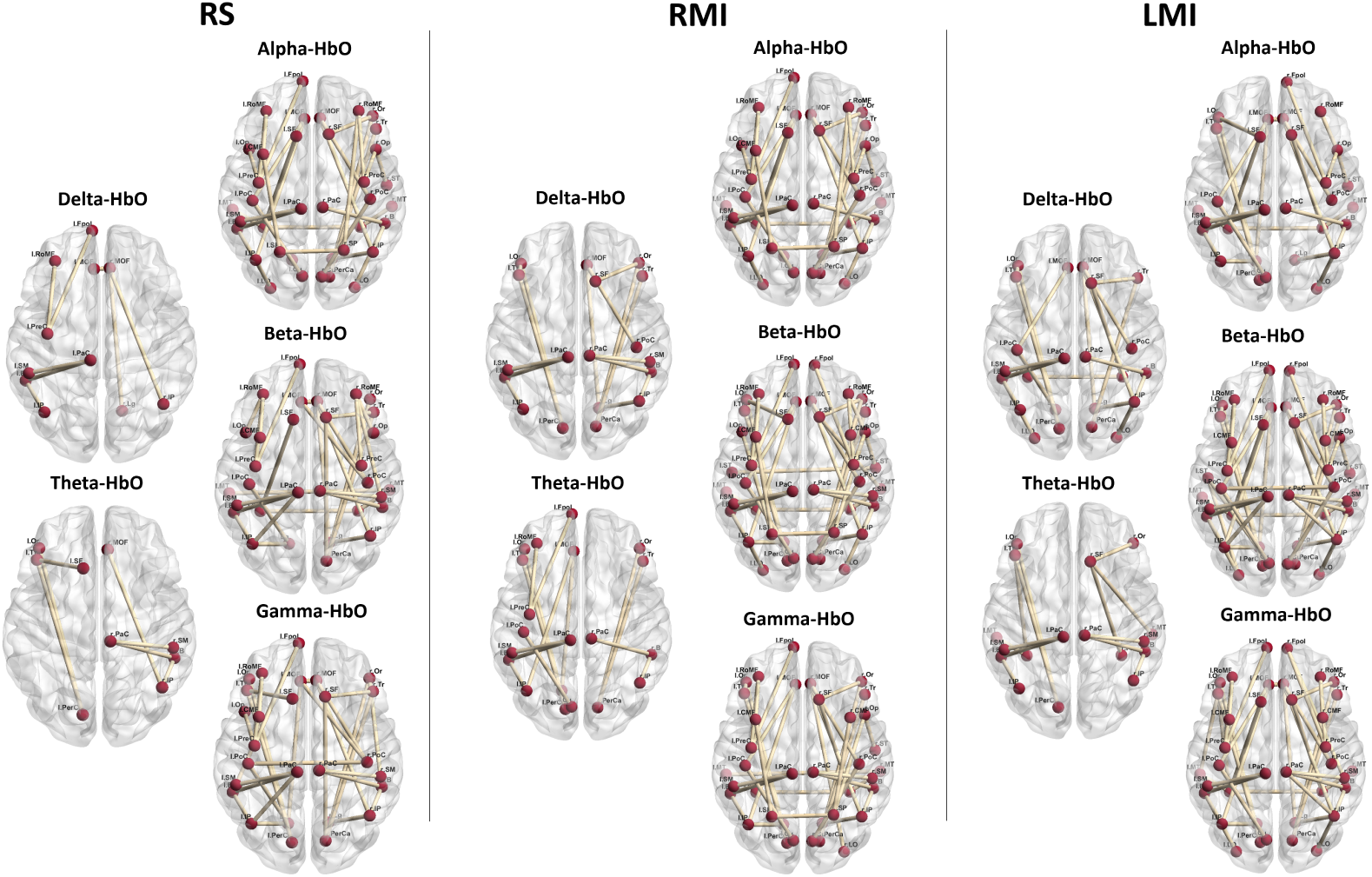
Stronger subnetwork in EEG bands across rest and task conditions compared to HbO networks.

**Figure 7:**
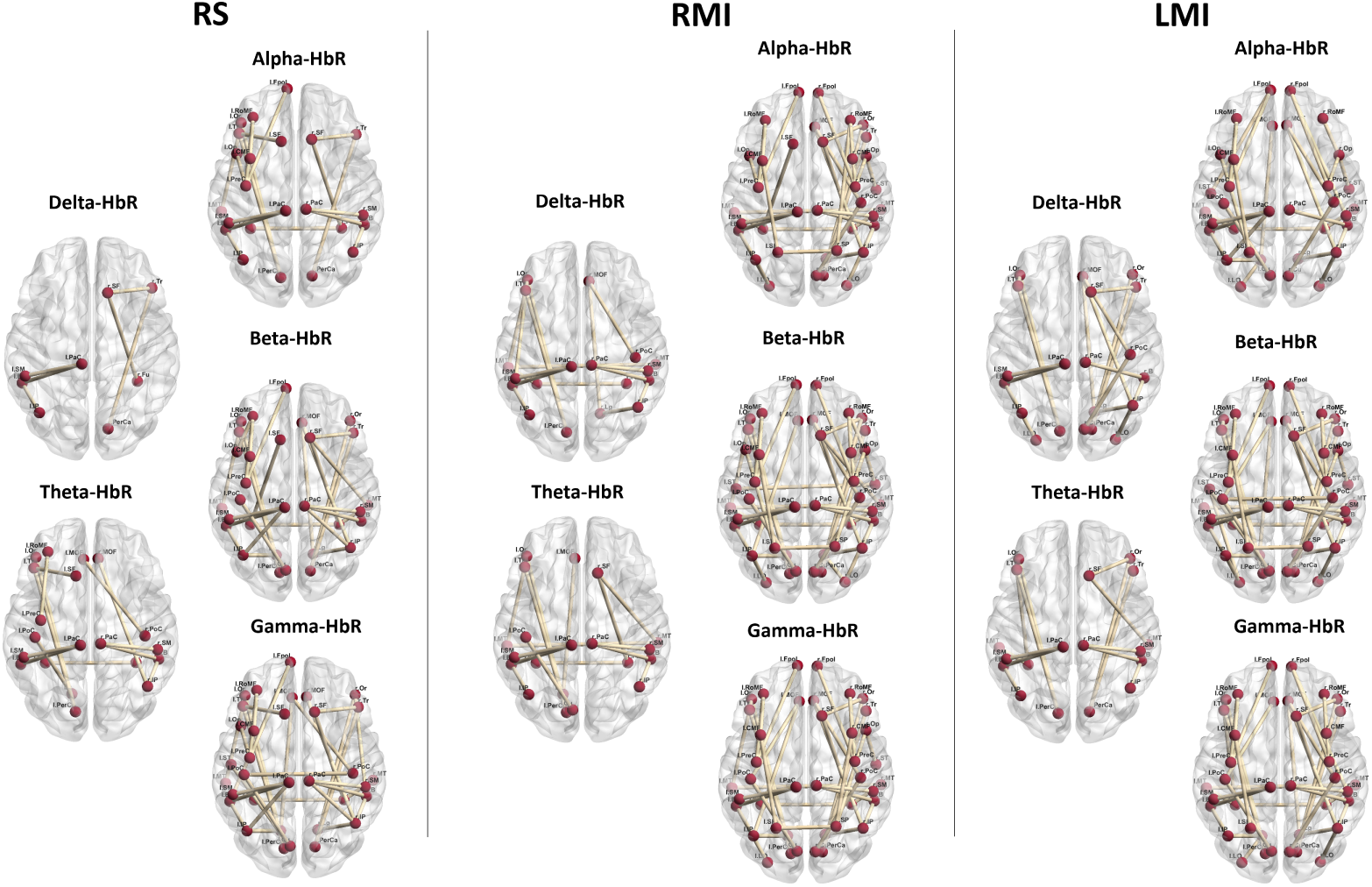
Stronger subnetwork in EEG bands across rest and task conditions compared to HbR networks.

During the task conditions (RMI and LMI), both showed a shared prominent subnetwork characterized by increased connectivity in *α*-HbO, *β*-HbO, and *γ*-HbO, as well as in *α*-HbR, *β*-HbR, and *γ*-HbR (Figures 8, 9). This subnetwork exhibited similar characteristics to those observed in the RS condition. However, for *θ*-HbO and *δ*-HbO, as well as *θ*-HbR and *δ*-HbR, distinct subnetworks with increased connectivity were identified. One of these subnetworks, consistent for both tasks, established connections between regions such as the bilateral inferior parietal (IP), bilateral supplementary motor area (PaC), and the right pars opercularis of the inferior frontal gyrus (Op). The other subnetwork, linked regions that included the bilateral medial frontal gyrus (mOF), right middle temporal gyrus (MT), and the right pars orbitalis of the inferior frontal gyrus (Or) in RMI. For LMI, it interconnected the bilateral medial frontal gyrus (mOF), right middle temporal gyrus (MT), and the right pars orbitalis (Or) and triangularis (Tr) of the inferior frontal gyrus. We then proceeded to examine the extent of similarity within each pair of resulting matrices across different frequency bands. Pearson correlation values (r) *>* 0.5 were chosen as the cutoff for determining similarity. Our findings revealed that only *θ*-HbO vs. *β*-HbO and *θ*-HbO vs. *δ*-HbO in the RS condition, and *θ*-HbR vs. *α*-HbR, *β*-HbR, and *γ*-HbR in the LMI condition, exhibited dissimilar subnetworks. In certain conditions and frequency bands, these FC matrices did not exhibit strong similarity, suggesting unique patterns in brain activity in these specific combinations of frequency bands and conditions.

**Figure 8:**
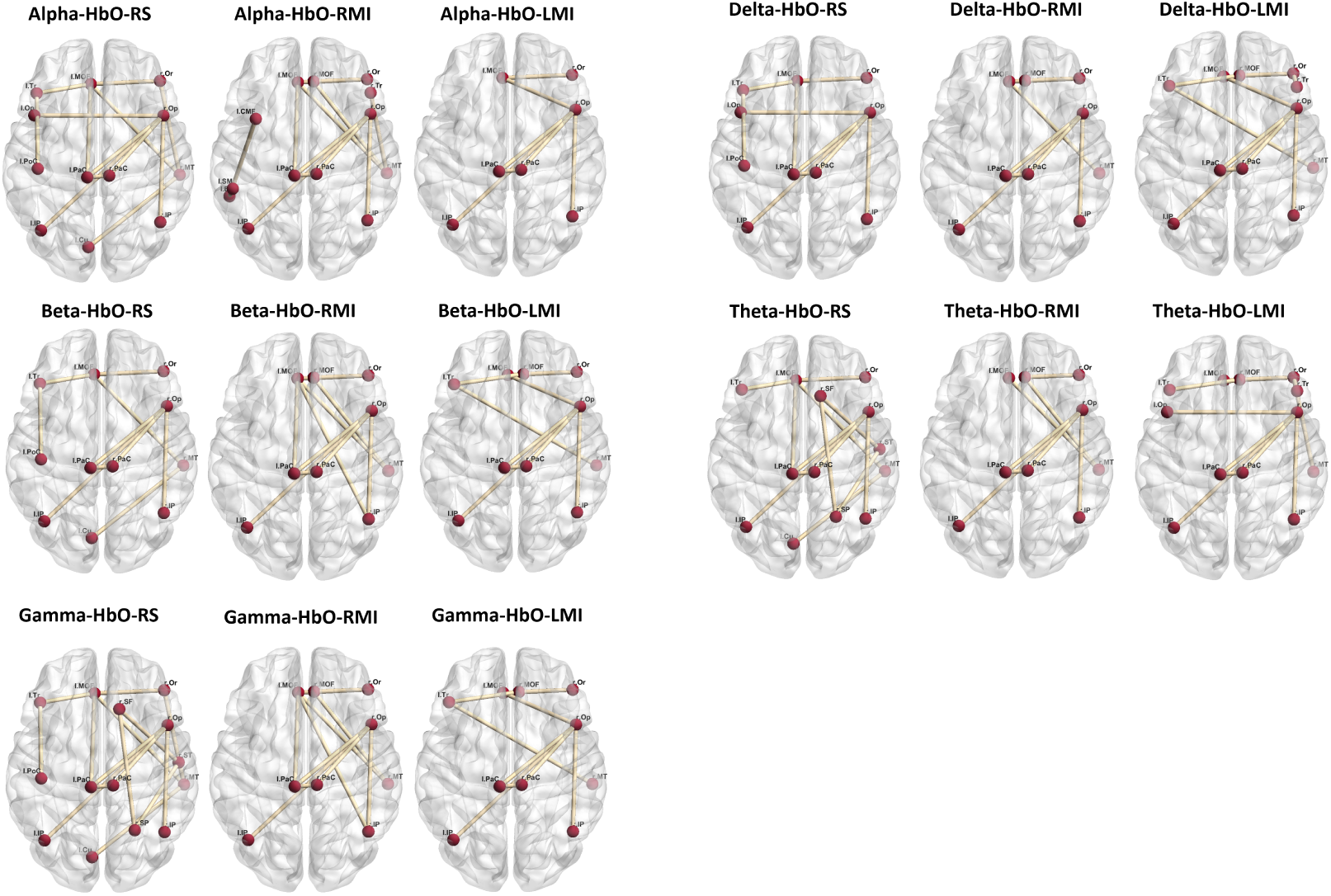
Stronger subnetwork in HbO networks across rest and task conditions compared to EEG bands.

**Figure 9:**
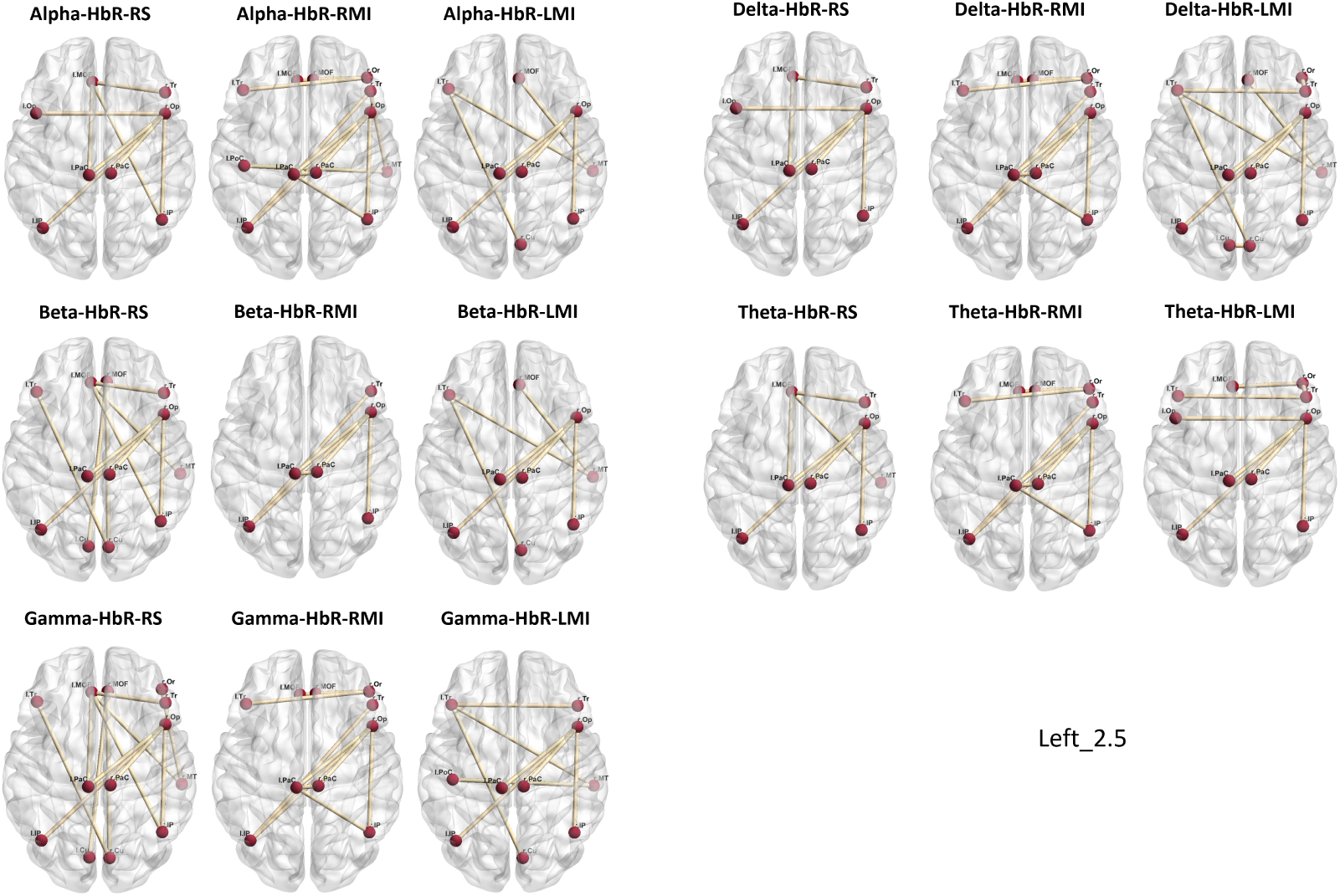
Stronger subnetwork in HbR networks across rest and task conditions compared to EEG bands.

However, when performing the right-tailed t-test, we pinpointed out subnetworks demonstrating increased functional connectivity within the EEG network relative to fNIRS. These subnetworks displayed intrahemispheric connections characterized by more clustered and distinct patterns across various frequency bands.

In the RS condition, we observed fewer differences between the two modalities than in tasks, particularly in *δ*-HbO and *θ*-HbO, where only two subnetworks were identified. These subnetworks connected regions such as the left pars orbitalis and pars triangularis of the inferior frontal gyrus (Or, Tr), the superior frontal gyrus (SF), and the pericalcarine cortex (PerCa) in one case, and the right inferior parietal (IP), medial orbitofrontal (mOF) right pericalcarine cortex (PerCa) in the other.

For *α*-HbO and *β*-HbO, we observed six subnetworks of higher connectivity, with two of them resembling the *δ*-HbO and *θ*-HbO patterns (Figure 6). Three shared intrahemispheric subnetworks exhibited more clustered patterns, connecting regions such as the pars opercularis of the inferior frontal gyrus (Op), pre-central gyrus (PreC), frontal cortex (Fpol), and rostral middle frontal cortex (RoMF) of the left hemisphere in the first subnetwork. The second subnetwork linked the same regions in the right hemisphere, while the third subnetwork connected temporo-parietal areas (B, IP, SM) and para-central regions (PaC). Similar patterns were identified in the task conditions, both for LMI and RMI. Furthermore, highly clustered intrahemispheric subnetworks were identified in *δ*-HbR, *θ*-HbR, *α*-HbR, *β*-HbR, and *γ*-HbR in all three conditions 7). The only non-significant result was observed in *θ*-HbR during the RS condition.

The similarity evaluation of these subnetworks across different frequency bands highlighted dissimilarity between *δ*-HbO vs. *θ*-HbO, *α*-HbO, and *γ*-HbO in RS, between *δ*-HbO vs. *β*-HbO, *α*-HbO, and *θ*-HbO in LMI, and between *δ*-HbO vs. *θ*-HbO, *α*-HbO, and *β*-HbO, as well as *θ*-HbO vs. *β*-HbO in RMI. Furthermore, subnetworks with r *<* 0.5 were predominantly identified during tasks in the case of *θ*-HbR vs. *α*-HbR, *β*-HbR, and *γ*-HbR for both LMI and RMI, while no such differences were observed in RS. In the case of *δ*-HbR vs. *α*-HbR, *β*-HbR, and *γ*-HbR in RMI, as well as *δ*-HbR vs. *α*-HbR and *β*-HbR in LMI. Dissimilarity was also evident in RS between *δ*-HbR vs. *β*-HbR and *γ*-HbR.

### 3.2. Multilayer networks

The last aim of this study was to investigate whether the integration of information from EEG and fNIRS provides a clearer picture of brain dynamics than the unimodal approach for inferring brain functions. To achieve this scope we chose a multilayer approach and compared the full-multilayers with the EEG- and fNIRS-multilayers by means of graph analysis. We chose the graph metrics, of Modularity (M), Global Efficiency (GE), and Path Length (PL) as measures of the effectiveness of the communication. By aggregating these measures across layers, the RS, RMI, and LMI conditions revealed distinctive modularity patterns within the Full-Multilayer network. Specifically, significantly higher modularity values were observed when comparing the Full-Multilayer to each of the EEG and the fNIRS networks in both RS and task conditions, indicating a more pronounced modular structure in the former (Figure 10). However, when comparing the Full-Multilayer network to the EEG-multilayer, in LMI and RMI conditions, its modularity values were only slightly higher, suggesting a degree of similarity between the two approaches. On the contrary, significantly lower M values were observed when contrasting the Full-Multilayer with the fNIRS-multilayer, as well as when comparing the EEG-multilayer with the fNIRS-multilayer.

**Figure 10:**
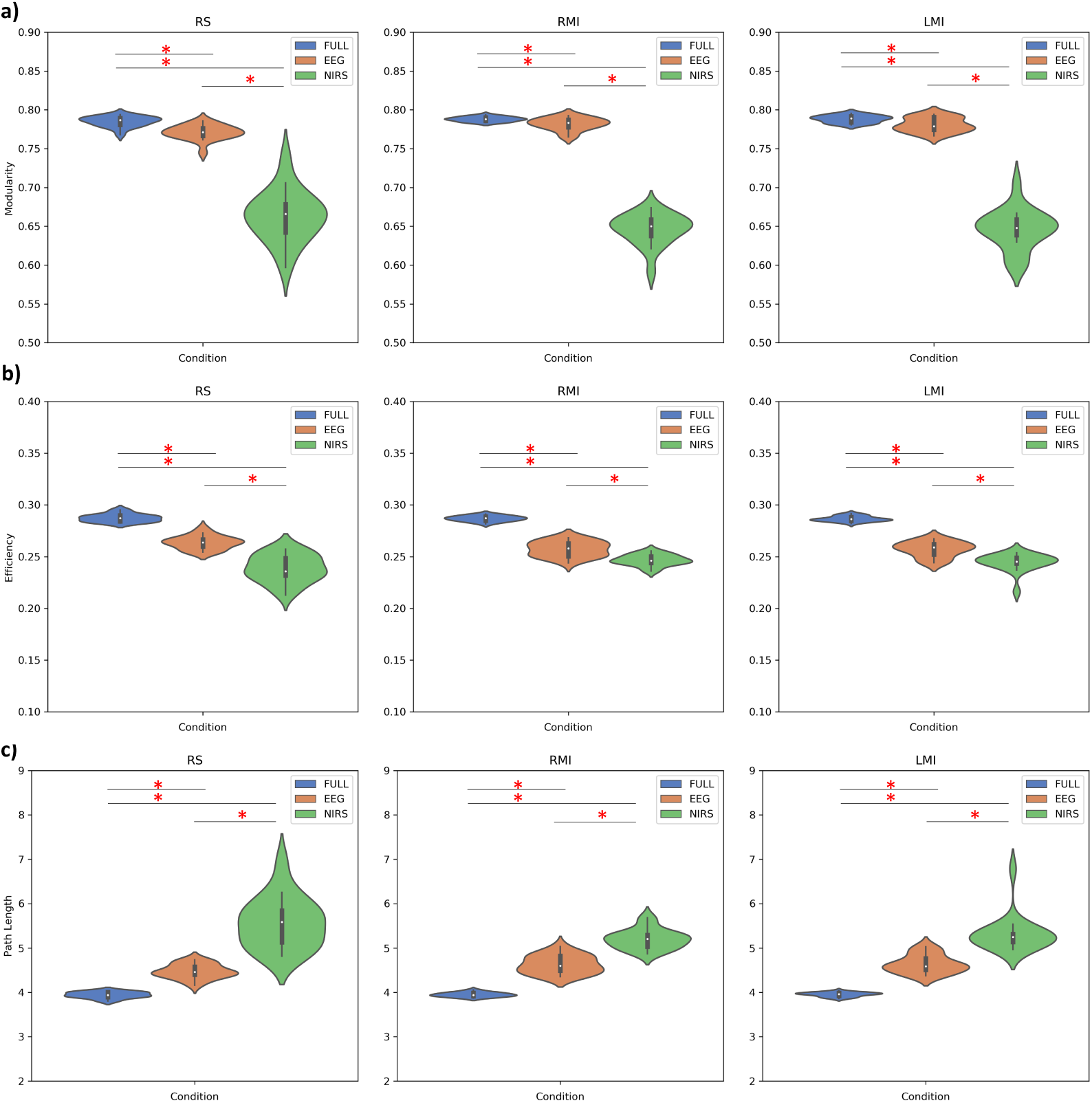
Comparison between the modularity (a) and the global efficiency (b) between multilayer and modality-specific networks.

These results suggested that the integrated approach excels in revealing modular structures in the brain that might otherwise remain obscured by unimodal techniques. Interestingly, while the multilayer approach surpassed fNIRS in terms of modularity, it also displayed slightly higher modularity values when compared to EEG. This suggests that the multilayer approach manages to capture more of the brain’s modular organization compared to fNIRS alone, but it may share some commonalities with EEG in this regard.

In all three conditions, RS, LMI, and RMI, the Full-Multilayer network exhibited significantly higher GE values when compared to both the EEG and fNIRS networks, indicating its superior capacity for global information transfer.

Furthermore, a statistically significant difference was also observed when comparing the EEG-multilayer with the fNIRS-multilayer in the three conditions, with the fNIRS network exhibiting lower GE values compared to the EEG one. This difference suggests that the EEG network may exhibit a more efficient global communication pattern compared to the fNIRS network in both RS and task conditions.

It’s noteworthy that the PL values showed an inverse trend. In all three conditions, RS, LMI, and RMI, significantly lower PL values were observed when comparing the Full-Multilayer network to both the EEG- and fNIRS-multilayer networks, suggesting more efficiency in terms of direct communication pathways between network nodes in the former. However, in RS, no significant differences were found between the EEG and fNIRS multilayer network, while in LMI and RMI, the fNIRS network exhibited significantly higher PL values compared to the EEG. This suggests that, for electrical brain activity, the neural information is routed via more globally optimal and shortest paths compared to hemodynamic activity during a cognitive task. Thus, providing faster and more direct information transfer.

Additionally, we investigated whether the integrated information in the full-multilayer provides complementary or redundant information compared to a single modality (EEG-multilayer or fNIRS-multilayer), in terms of spatial distribution of the active brain regions. We conducted this assessment by computing the Dice coefficient based on EC values to take into account the spacial information. The high similarity suggests that the integrated information has a similar effect on capturing network structure compared to a single modality, indicating redundancy. Conversely, low similarity suggests that the integration of different modalities leads to differences in network structure, emphasizing the presence of complementary information.

We observed, in the RS condition, a Dice coefficient of 0.91 when comparing the EC values of the Full-Multilayer with those of the EEG-Multilayer. This means that many brain regions exhibit similar EC values during RS, indicating redundancy between the two approaches. This implies that the EEG modality is suitable to characterize brain patterns in RS when used in isolation. In contrast, the comparison of Full-Multilayer EC values with fNIRS-Multilayer EC values yielded a Dice coefficient of 0.68. This suggests that the fNIRS approach captures somewhat different patterns of brain centrality during RS, implying that a multimodal approach can provide complementarity when different sources of information are integrated.

Similar trends were shown during the task conditions. For both LMI and RMI, a higher coefficient of similarity was highlighted for EEG (Dice = 0.88 in RMI, Dice = 0.84 in LMI) indicating substantial similarity with full-multilayer, compared to fNIRS which showed lower values (Dice = 0.72 in RMI, Dice = 0.63 in LMI), suggesting some differences with the integrated approach (Table 2). This indicated a certain level of redundancy and complementarity between the multimodal and the unimodal approach that is dependent on the modality and on the specific brain state, but also between the different types of tasks or the cognitive demand during the tasks.

**Table 2:**
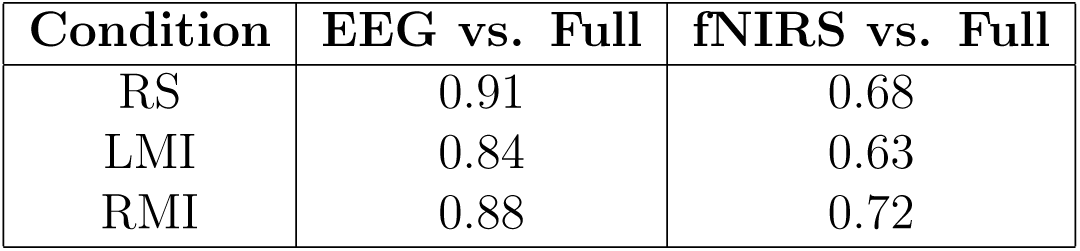
Dice Coefficients for the three conditions (RS, LMI, RMI)

**Table 3:**
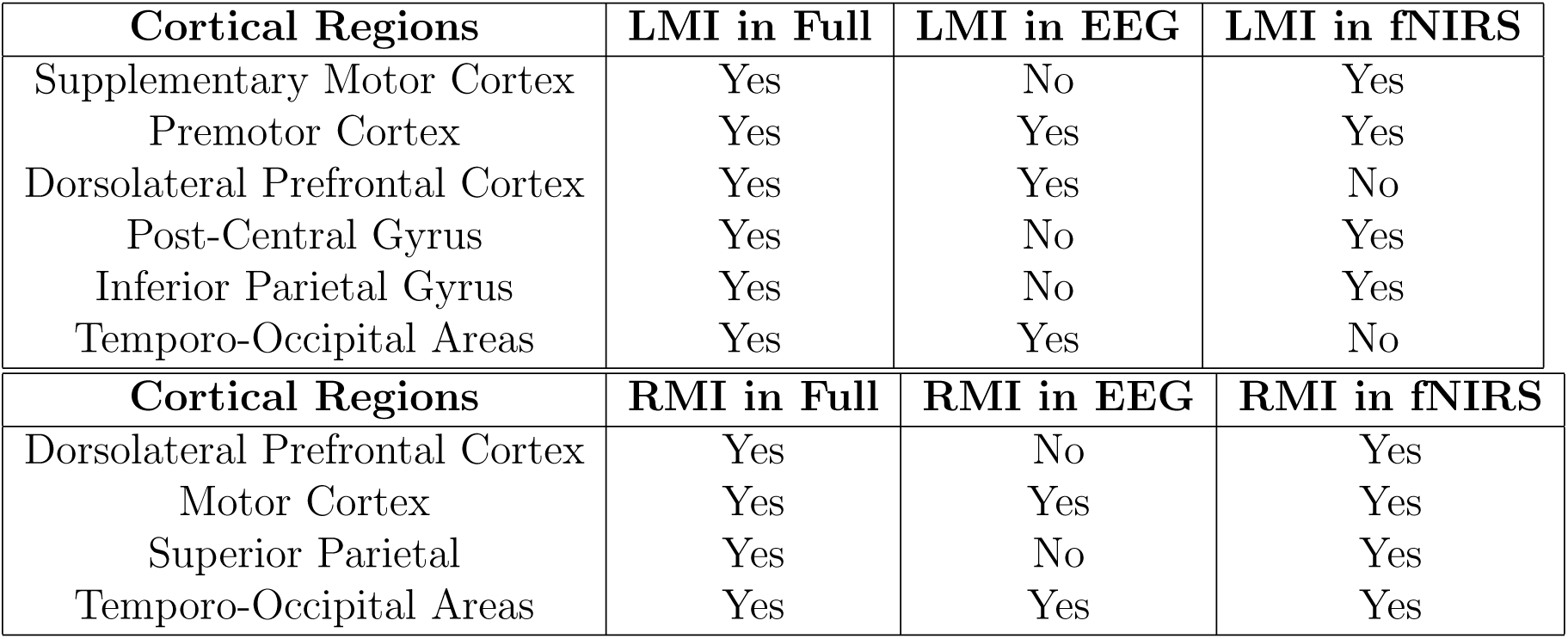
Active regions during LMI and RMI.

Then, we systematically computed differences in each task condition relative to the RS as a baseline, to capture variations in brain activity during LMI and RMI for the Full-Multilayer, EEG-Multilayer, and fNIRS-Multilayer, separately. Subsequently, we mapped the contrasts on the chosen cortical regions linked to the MI for all three multilayers.

We found that during LMI, in the full-multilayer, 12 out of 26 cortical regions involved in MI were activated, including the supplementary motor cortex (PaC), premotor cortex (SF), dorsolateral prefrontal cortex (RoMF, CMF), post-central gyrus (PoC), inferior parietal gyrus (IP), and temporo-occipital associative areas (Fu, Cu, MT, ST).

In the EEG-multilayer, 8 out of 26 cortical regions including regions like the dorsolateral prefrontal cortex (CMF), premotor cortex (SF), the postcentral gyrus (PoC), and temporo-occipital association areas (Fu, Cu, MT, ST).

In the fNIRS-multilayer, 5 out of 26 regions comprising areas like the premotor cortex (SF), pre-central gyrus (PreC), and the supplementary motor area (PaC), superior parietal gyrus (SP), and superior temporal gyrus (ST). During RMI, in the full-multilayer 6 out of 26 cortical regions involved in MI were activated, including the dorsolateral prefrontal cortex (RoMF), motor cortex (PreC), superior parietal (SP), and temporo-occipital associative areas (Fu, Cu, ST).

In the EEG-multilayer, 6 out of 26 regions comprising the dorsolateral premotor cortex (CMF), motor cortex (PreC), and temporo-occipital associative areas (MT, Fu, Cu, ST).

In the fNIRS-multilayer, 12 out of 26 regions including the somatosensory and motor cortex (PoC, PreC), dorsolateral prefrontal cortex (CMF, RoMF), supplementary motor area (PaC), premotor cortex (SF), superior and inferior parietal gyrus (SP, IP), and temporo-associative areas (ST, MT).

These findings are summarized in Table tab:cortical-regions and indicate differences and similarities of activated brain regions during the task conditions across modalities.

## 4. Discussion

In this study, our primary focus was to examine the correspondence between fNIRS functional connectivity and EEG functional connectivity. Our emphasis was on discerning both shared characteristics and divergences in network topology between these two modalities across frequency bands through graph analysis. Additionally, we investigated how the interplay between electrophysiological and hemodynamic activities varies between the resting state (RS) and task-related conditions (left and right motor imagery task namely LMI and RMI respectively).

We observed a small-world propensity for both modalities in the three conditions (RS, LMI, and RMI). The small-world structure of a network characterizes its efficiency and cost-effectiveness. Our findings agree with previous results, supporting the assumption that small-worldness is a universal principle for the functional wiring of the human brain regardless of the distinct mechanisms of different imaging techniques [3]. However, the brain supports both segregated and distributed information processing, a key for cognitive processing, which means that localized activity in specialized regions is spread by coherent oscillations in large-scale distributed systems [44]. The global clustering coefficient (GCC) measures the clustering ability within the network, while the characteristic path length (PL) assesses how efficiently information travels within the network. We showed that all EEG frequency bands had significantly higher GCCs than HbO and HbR, while all EEG and fNIRS networks exhibited PL values of approximately 1. This indicates that the electrical networks have a better ability to form clusters or subgroups of interconnected nodes in the brain, but also that both EEG and fNIRS networks exhibit an efficient information transfer across the network, thus resulting in a small-world topology. This topology was observed for both the RS and the task conditions. Notably, during the RS, the *α* band exhibited higher clustering, aligning with its predominance during periods of rest and in agreement with previous studies. In contrast, during the tasks, we observed increased GCC in the *β* and *γ* bands, frequencies closely associated with motor imagery tasks [45], [35], [20], [40].

Moreover, the findings are consistent with the understanding that different brain networks are selectively engaged with specific frequency bands. For instance, the dorsal attentional network (DAN) and somatomotor network (SMN) are primarily associated with alpha and beta oscillations, and the ventral attentional network (VAN) is linked to gamma oscillations [38].

Our observations revealed higher global efficiency (GE) for the EEG network compared to the fNIRS (HbO) network only during RS and only in *θ* and *β* frequency bands, while in *θ* band as compared to HbR. This suggests that during RS, EEG shows higher information exchange capacity in specific frequency bands compared to fNIRS. On the contrary, in task conditions, when compared to HbO, no differences were pointed out, suggesting that the efficiency of distributed information transfer in the brain network between the two modalities is more similar when increasing the synchronization of neuronal oscillations for sustained brain activation, such as during a cognitive task. In fact, functional similarity between neighboring units minimizes the required wiring and may contribute to the high efficiency of information transfer [8].

These differences and similarities between the two modalities could be related to the different sensitivities and the temporal dynamics of these modalities to neural activity. The highest temporal resolution of EEG allows to capture of faster changes in neural activity, thus providing a more precise estimation of the timing of information transfer between brain regions in RS, which could contribute to higher global efficiency. On the contrary, fNIRS provides insights into the slower hemodynamic responses associated with longer-lasting and sustained neural processes, comprising cognitive tasks that involve extensive coordination across various brain regions [36].

Additionally, the variation observed in PL (lower values) and GCC (higher values) within the EEG network for almost all frequency bands compared to both HbO and HbR, suggests that, electrical networks have a better ability to form clusters of interconnected nodes in the brain and that the neural information is routed via more globally optimal and shortest paths compared to hemodynamic activity. This provides faster and more direct information transfer [8]. These findings are consistent for both RS and task conditions, highlighting the complex interplay of frequency bands and brain state in shaping the brain’s functional connectivity and network structure [45].

Cognitive states involve the engagement of different oscillatory frequencies within cortical networks, often lasting mere milliseconds during resting states [7], [23] and those states often last only a few ms. It is widely accepted that the short- and long-range organizations of neural oscillations from different frequencies depend on how those oscillations work together and that more than one EEG frequency band has been associated with the same resting state network [38], [45], [20]. Therefore, there is an advantage of EEG in capturing these transient neuronal activations. In contrast, fNIRS is limited in its ability to distinguish these rapidly changing neural responses, given that the hemodynamic response is delayed by several seconds, thus reflecting the sum of several oscillatory network configurations. [38],

The edge-wise analysis highlighted a large subnetwork with increased connectivity for the fNIRS network over EEG, exhibiting preferentially interhemispheric and homotopic connections between various cortical regions. Also, EEG networks demonstrated increased functional connectivity over fNIRS networks in intrahemispheric connections that are characterized by more clustered and distinct patterns across various frequency bands. The findings may imply that the generation of homologous connectivity is through direct structural connection, while intrahemispheric connectivity may reflect the synchronization of transient neural activation among distant cortical regions. Several studies suggested that oscillations at higher and lower frequencies may support short- and long-range connectivity patterns, respectively, and that slower oscillations are better suited than faster oscillations for supporting long-range connectivity in the brain, which in turn promotes high-order brain functions [12] [28] [32]. Indeed, we observed fewer subnetwork differences between the two modalities in RS condition than in tasks, particularly in slower frequency (*δ*-HbO and *θ*-HbO), while during cognitive tasks highly clustered intrahemispheric subnetworks emerged for all the frequency bands.

When assessing the similarities of these patterns across frequency bands, for the networks favoring fNIRS over EEG, we observed dissimilar subnetworks only in the RS condition for *θ*-HbO vs. *δ*-HbO and *β*-HbO, and only in LMI conditions for *θ*-HbR vs. *α*-HbR, *β*-HbR, and *γ*-HbR. These findings can be explained by the different sensitivities and temporal resolution of the two modalities and by the coupling of electrical and hemodynamic activity governed by the neurovascular mechanism. Transient increases in neuronal synchronization in the alpha, beta, or gamma frequencies are typically followed by a decline in the oxygenated hemoglobin concentration (because of its usage), thus correlating with lower metabolic demand [41], [30].

On the contrary, when investigating the similarity of these subnetworks across different frequency bands, for the networks favoring EEG over fNIRS, differences were highlighted between lower (*δ*-HbO, *θ*-HbO, and *δ*-HbR, *θ*-HbR) vs. higher frequency bands (*α*-HbO, *β*-HbO, and *γ*-HbO, and *α*-HbR, *β*-HbR, and *γ*-HbR) in both RS and tasks. The dissimilarity between specific frequency bands in the RS condition suggests that even during a supposed state of rest, the brain exhibits variability in activity patterns. This variability might be due to factors such as internal thought processes, background sensory input, or fluctuations in attention [38]. In contrast, during tasks observed dissimilarity can be attributed to the cognitive demands and specific neural processes associated with these tasks. Task-related dissimilarity could be due to the dynamic reconfiguration of neural networks to adapt to the cognitive demands of the tasks and may be indicative of how different cognitive functions engage distinct brain regions or networks [7], [45]. It was suggested that the preferential neural oscillation for network connectivity depends on the spatial distribution of the RSN nodes and structural connections in the brain [19], [54]. However, future studies may provide novel insights into the possible relationships between network topology and preferential oscillations.

In the final objective of our study, we employed a multilayer approach to examine whether it outperforms the unimodal approach in characterizing the topology of the brain network, thus providing a clearer picture of brain functioning. Additionally, we probed whether the integrated information in the full-multilayer provides complementary or redundant information compared to a single modality. We compared the full-multilayers with the EEG- and fNIRS-multilayers by means of graph analysis. We chose the graph metrics, of Modularity (M), Global Efficiency (GE), and Path Length (PL) as measures of the effectiveness of the communication. We found significantly higher M values when comparing the Full-Multilayer to each of the EEG and the fNIRS networks in both RS and task conditions, indicating a more pronounced modular structure in the Full-Multilayer. These results suggested that the integrated approach excels in revealing modular structures in the brain that might otherwise remain obscured by unimodal techniques. Interestingly, while the full-multilayer surpassed the fNIRS-multilayer in terms of modularity, it also displayed slightly higher M values when compared to the EEG-multilayer. This means that the full-multilayer captures more of the brain’s modular organization compared to the fNIRS network alone, but it may share some commonalities with the EEG network in this regard. Beyond M, we also examined GE and PL within the networks. The Full-Multilayer consistently outperformed both the EEG-multilayer and the fNIRS-multilayer in terms of GE in all three conditions (RS, LMI, RMI), indicating its superior capacity for global information transfer. Notably, PL values showed an inverse trend. The Full-Multilayer consistently exhibited significantly lower PL values compared to both the EEG- and fNIRS-multilayer, underscoring its efficiency in terms of direct communication pathways between network nodes. Several other studies have similarly determined that a multilayer network model contains non-trivial information that cannot be retrieved by considering individual layers independently [4], [13], [14]. The primary rationale behind this phenomenon is predominantly conceptual and organizational. Vector labels, which encompass attributes assigned to nodes or layers in the multilayer network, such as measures of connectivity strength, gene expression profiles, or neurotransmitter levels in specific brain regions, provide a means of identifying meaningful subgroups within the comprehensive dataset. The analytic power of a multilayer network comes from comparing statistics across these subsets. The multilayer networks and network statistics are designed to operate across layers, thus the full multidimensional information can be retained, allowing for a richer analysis that could not be obtained from the aggregated single-layer network [60].

Additionally, we investigated whether the integrated information in the full-multilayer provides complementary or redundant information compared to a single modality by means of the Dice coefficient of similarity.

The results reveal differences between the multimodal and the unimodal approach. Specifically, during the RS, the Dice coefficient of 0.91 between the EC values obtained from the Full-Multilayer and those from the EEG-Multilayer indicates that many brain regions exhibit similar EC values, suggesting redundancy. This implies that EEG, when used in isolation, is suitable for characterizing brain patterns during the resting state. Conversely, the Dice coefficient of 0.68 between the EC values obtained from the Full-Multilayer and those from the fNIRS-Multilayer suggests that the fNIRS network captures slightly different patterns of brain centrality. In this case, fNIRS appears to add information that is not present in EEG, suggesting that a multimodal approach can provide complementarity when different sources of information are integrated, thus enhancing the understanding of brain centrality during the resting state. Similar trends were observed during tasks involving motor imagery. For both LMI and RMI, EEG exhibited a higher Dice coefficient compared to fNIRS when compared to the Full-Multilayer. Overall, these results indicate the presence of some level of overlap and complementarity between the multimodal and unimodal approaches, which is influenced by the detection modality, specific brain state, and the type of activity performed. These findings underscore the importance of considering a multimodal approach to gain a more comprehensive understanding of brain patterns in different contexts [43], [57], [25].

To gain further insights, we systematically computed differences in each task condition relative to the RS as a baseline, to capture variations in brain activity during LMI and RMI for the Full-Multilayer, EEG-Multilayer, and fNIRS-Multilayer, separately. We mapped these contrasts on the chosen cortical regions linked to the MI for all three multilayers, to see if this approach outperformed the unimodality in capturing the centrality of the network.

We found that, for both LMI and RMI, the full-multilayer identified a higher number of central nodes in the network than the EEG-multilayer, while compared to the fNIRS-multilayer only in LMI. The nodes identified and shared with full-multilayer were different from the EEG and fNIRS network. Specifically, fronto-posterior regions were detected by EEG-multilayer comprising the dorsolateral prefrontal cortex and temporo-occipital associative area, while fNIRS-multilayer captured brain activity in inferior parietal gyrus, post-central gyrus, and supplementary motor area. Remarkably, the premotor cortex was detected by both.

We found that during LMI, 3 out of 6 cortical regions identified by fullmultilayer, were captured by EEG-multilayer: DLPC, pre-motor cortex, and tempo-occipital areas. On the contrary, 4 out of 6 regions identified by fullmultilayer involved, were captured by fNIRS-multilayer: SMA, pre-motor cortex, post-central gyrus, and inferior-parietal gyrus. Only one region overlaps between EEG- and fNIRS-multilayer. In RMI, 2 out of 4 regions identified by full-multilayer, were also detected by EEG: motor cortex and temporo-occipital areas, while for fNIRS 4 out of 4: DLPC, motor cortex, superior parietal gyrus, and temporo-occipital area.

Our results suggest that while there is a degree of overlap in activated regions across modalities, there are also unique contributions from each modality. The full-multilayer approach, which integrates EEG and fNIRS data, captured a higher number of nodes activated during both LMI and RMI, than the unimodal approach. This indicates the potential for enhanced characterization of brain functions when different techniques are combined. For example, Battiston and colleagues (2017) [4] employed a two-layer multiplex network combining fMRI- and DTI-based connectivity networks to explore the recurrence of newly defined multilayer motifs. Their findings revealed significant variations in these motifs that were over-represented in the human brain, distinguishing them from previous findings solely based on structural connectivity. Similarly, in the work of Crofts and colleagues (2016) [11], a novel measure of structure-function clustering enabled them to probe functional connections that deviate from the underlying cortical structure.

Our findings agree with previous studies demonstrating that a multilayer representation of a human brain network can provide new insight into pathology. For example, greater classification accuracy in distinguishing between healthy and schizophrenic patients was reported when compared to single-layer networks [16], or improvement in the discrimination between Alzheimer’s disease and healthy populations [9].

Moreover, we observed that specific cortical regions were consistently activated across different modalities. This suggests that EEG and fNIRS, to some extent, can provide similar information about the neural processes occurring in certain cortical regions. This is advantageous when developing classification algorithms [63] or applications for brain-computer interfaces [37].

## 5. Conclusion, Limitations and Future Directions

Our study has successfully achieved its aim of comparing EEG and fNIRS and assessing the effectiveness of the multilayer approach. The results highlight differences in how EEG and fNIRS capture brain network topology in RS and tasks. EEG excels in capturing rapid changes during the resting state, while fNIRS provides insights into slower processes during cognitive tasks. These variations are linked to their distinct sensitivities and temporal dynamics. The specific neural oscillations involved might depend on the brain’s structure and resting-state network distribution. Future studies may provide novel insights into the possible relationships. From a multimodal perspective, we demonstrated that the multilayer approach offers advantages in revealing modular organization and improving global information transfer in the brain network. These findings have implications for neuroimaging research, highlighting the potential benefits of integrating multiple modalities for a more comprehensive understanding of brain function and organization. There are a number of limitations to the work presented here. A small sample size can limit the ability to draw robust conclusions. Despite meticulous preprocessing, data quality can vary between subjects and modalities, affecting the reliability of connectivity measures. EEG and fNIRS had limited spatial coverage of the sensors on the scalp resulting in reduced brain regions with respect to the original Atlas. Motor imagery tasks exhibit high variability across subjects because of the difficulty of the task. We did not consider within-subject analysis which may account for this variability.

From a more theoretical point of view, it is possible that the multilayer model itself could be expanded. For example, this framework allows for the integration of multiple modalities (e.g. EEG/fNIRS) and structural networks. Previous studies have demonstrated that temporal networks exhibit distinctive phases of segregation and integration. Future investigations in dynamic multilayer analysis could delve into this phenomenon and explore how multilayer functional hubs facilitate the coordination of dynamical processing. Finally, a practical implication of the current findings is that in future studies, researchers should carefully weigh the decision to employ a specific neuroimaging technique, taking into account its respective strengths and limitations and how this choice may impact their research outcomes.

## Acknowledgements

This work was funded by the European Union’s Horizon 2020 research and innovation program grant Sano no 857533, and the International Research Agendas program of the Foundation for Polish Science, co-financed by the European Union under the European Regional Development Fund.

## Declaration of Generative AI and AI assisted technologies in the writing process

During the preparation of this work, the author(s) used ChatGPT for grammar and spell check. After using this tool, the author(s) reviewed and edited the content as needed and take(s) full responsibility for the content of the publication.

## Data availability

The dataset used in this study is available at: https://doc.ml.tuberlin.de/hBCI/contactthanks.php All the figures in the document need color for printing.

## Author contributions

Rosmary Blanco: Conceptualization, Methodology, Data Analysis, Writing-Original draft preparation, Writing-Reviewing and Editing, Visualization. Cemal Koba: Writing-Reviewing and Editing. Alessandro Crimi: Supervision

## References

[1] Abdalmalak, A., Novi, S.L., Kazazian, K., Norton, L., Benaglia, T., Slessarev, M., Debicki, D.B., Lawrence, K.S., Mesquita, R.C. and Owen, A.M., 2022. Effects of systemic physiology on mapping resting-state networks using functional near-infrared spectroscopy. Frontiers in neuroscience, 16, p.803297.

[2] Angermann, A., Beuschel, M., Rau, M. and Wohlfarth, U., 2004. Matlab-Simulink-Stateflow. Grundlagen, Toolboxen.

[3] Bassett, D.S. and Bullmore, E.T., 2017. Small-world brain networks revisited. The Neuroscientist, 23(5), pp.499–516.

[4] Battiston, F., Nicosia, V. and Latora, V., 2014. Structural measures for multiplex networks. Physical Review E, 89(3), p.032804.

[5] Blanco, R., Koba, C., Crimi, A., 2023 ”Resting State Brain Connectivity Analysis from EEG and FNIRS Signals” In: Miky̌ska, J., de Mulatier, C., Paszynski, M., Krzhizhanovskaya, V.V., Dongarra, J.J., Sloot, P.M. (eds) Computational Science – ICCS 2023. ICCS 2023. Lecture Notes in Computer Science, vol 14074. Springer, Cham. 10.1007/978-3-031-36021-358

[6] Breedt, L.C., Santos, F.A., Hillebrand, A., Reneman, L., van Rootselaar, A.F., Schoonheim, M.M., Stam, C.J., Ticheler, A., Tijms, B.M., Veltman, D.J. and Vriend, C., 2023. Multimodal multilayer network centrality relates to executive functioning. Network Neuroscience, 7(1), pp.299–321

[7] Brookes, M.J., Woolrich, M., Luckhoo, H., Price, D., Hale, J.R., Stephenson, M.C., Barnes, G.R., Smith, S.M. and Morris, P.G., 2011. Investigating the electrophysiological basis of resting state networks using magnetoencephalography. Proceedings of the National Academy of Sciences, 108(40), pp.16783–16788.

[8] Bullmore, E. and Sporns, O., 2009. Complex brain networks: graph theoretical analysis of structural and functional systems. Nature reviews neuroscience, 10(3), pp.186–198.

[9] Cai, L., Wei, X., Liu, J., Zhu, L., Wang, J., Deng, B., Yu, H. and Wang, R., 2020. Functional integration and segregation in multiplex brain networks for Alzheimer’s disease. Frontiers in Neuroscience, 14, p.51.

[10] Cai, Z., Machado, A., Chowdhury, R.A., Spilkin, A., Vincent, T., Aydin, Ü., Pellegrino, G., Lina, J.M. and Grova, C., 2021. Diffuse optical reconstructions of fNIRS data using Maximum Entropy on the Mean. bioRxiv, pp.2021–02.

[11] Crofts, J.J., Forrester, M. and O’Dea, R.D., 2016. Structure-function clustering in multiplex brain networks. Europhysics Letters, 116(1), p.18003.

[12] Da Silva, F.L., 2013. EEG and MEG: relevance to neuroscience. Neuron, 80(5), pp.1112–1128.

[13] De Domenico, M., Solé-Ribalta, A., Cozzo, E., Kivelä, M., Moreno, Y., Porter, M.A., Gómez, S. and Arenas, A., 2013. Mathematical formulation of multilayer networks. Physical Review X, 3(4), p.041022

[14] De Domenico, M., Nicosia, V., Arenas, A. and Latora, V., 2014. Layer aggregation and reducibility of multilayer interconnected networks. arXiv preprint arXiv:1405.0425.

[15] De Domenico, M., Porter, M.A. and Arenas, A., 2015. MuxViz: a tool for multilayer analysis and visualization of networks. Journal of Complex Networks, 3(2), pp.159–176.

[16] De Domenico, M., Sasai, S. and Arenas, A., 2016. Mapping multiplex hubs in human functional brain networks. Frontiers in neuroscience, 10, p.326.

[17] Desikan, R.S., Śegonne, F., Fischl, B., Quinn, B.T., Dickerson, B.C., Blacker, D., Buckner, R.L., Dale, A.M., Maguire, R.P., Hyman, B.T. and Albert, M.S., 2006. An automated labeling system for subdividing the human cerebral cortex on MRI scans into gyral based regions of interest. Neuroimage, 31(3), pp.968–980.

[18] Esfahlani, F.Z. and Sayama, H., 2018. A percolation-based thresholding method with applications in functional connectivity analysis. In Complex Networks IX: Proceedings of the 9th Conference on Complex Networks CompleNet 2018 9 (pp. 221–231). Springer International Publishing.

[19] Fukushima, M. and Sporns, O., 2020. Structural determinants of dynamic fluctuations between segregation and integration on the human connectome. Communications biology, 3(1), p.606.

[20] Ganzetti, M. and Mantini, D., 2013. Functional connectivity and oscillatory neuronal activity in the resting human brain. Neuroscience, 240, pp.297–309.

[21] Gramfort, A., Papadopoulo, T., Olivi, E. and Clerc, M., 2010. OpenMEEG: opensource software for quasistatic bioelectromagnetics. Biomedical engineering online, 9, pp.1–20.

[22] Gramfort, A., Luessi, M., Larson, E., Engemann, D.A., Strohmeier, D., Brodbeck, C., Goj, R., Jas, M., Brooks, T., Parkkonen, L. and Hämäläinen, M., 2013. MEG and EEG data analysis with MNE-Python. Frontiers in neuroscience, p.267.

[23] Hacker, C.D., Snyder, A.Z., Pahwa, M., Corbetta, M. and Leuthardt, E.C., 2017. Frequency-specific electrophysiologic correlates of resting state fMRI networks. Neuroimage, 149, pp.446–457.

[24] Hagberg, A., Swart, P. and S Chult, D., 2008. Exploring network structure, dynamics, and function using NetworkX (No. LA-UR-08-05495; LA-UR-08-5495). Los Alamos National Lab.(LANL), Los Alamos, NM (United States).

[25] Hammoud, Z. and Kramer, F., 2020. Multilayer networks: aspects, implementations, and application in biomedicine. Big Data Analytics, 5(1), p.2.

[26] He, B. and Liu, Z., 2008. Multimodal functional neuroimaging: integrating functional MRI and EEG/MEG. IEEE reviews in biomedical engineering, 1, pp.23–40.

[27] Hiyoshi, H. and Sugihara, K., 2000, May. Voronoi-based interpolation with higher continuity. In Proceedings of the sixteenth annual symposium on Computational geometry (pp. 242–250).

[28] Jones, S.R., Pinto, D.J., Kaper, T.J. and Kopell, N., 2000. Alphafrequency rhythms desynchronize over long cortical distances: a modeling study. Journal of computational neuroscience, 9, pp.271–291.

[29] Kivelä, M., Arenas, A., Barthelemy, M., Gleeson, J.P., Moreno, Y. and Porter, M.A., 2014. Multilayer networks. Journal of complex networks, 2(3), pp.203–271.

[30] Koch, S.P., Koendgen, S., Bourayou, R., Steinbrink, J. and Obrig, H., 2008. Individual alpha-frequency correlates with amplitude of visual evoked potential and hemodynamic response. Neuroimage, 41(2), pp.233–242.

[31] Kohno, S., Miyai, I., Seiyama, A., Oda, I., Ishikawa, A., Tsuneishi, S., Amita, T. and Shimizu, K., 2007. Removal of the skin blood flow artifact in functional near-infrared spectroscopic imaging data through independent component analysis. Journal of biomedical optics, 12(6), pp.062111–062111.

[32] Kopell, N., 2000. We got rhythm: Dynamical systems of the nervous system. Notices of the AMS, 47(1), pp.6–16.

[33] Kruskal, J.B., 1956. On the shortest spanning subtree of a graph and the traveling salesman problem. Proceedings of the American Mathematical society, 7(1), pp.48–50.

[34] Krylova, M.A., Izyurov, I.V., Gerasimenko, N.Y., Slavytskaya, A.V. and Mikhailova, E.S., 2016. Human brain networks for visual spatial orientations processing. Fechner Day 2016, p.85.

[35] Leeuwis, N., Yoon, S. and Alimardani, M., 2021. Functional connectivity analysis in motor-imagery brain computer interfaces. Frontiers in Human Neuroscience, 15, p.732946.

[36] Li, R., Yang, D., Fang, F., Hong, K.S., Reiss, A.L. and Zhang, Y., 2022. Concurrent fNIRS and EEG for brain function investigation: a systematic, methodology-focused review. Sensors, 22(15), p.5865.

[37] Llorente, D., Ballesteros, M., Cruz-Ortiz, D., Salgado, I. and Chairez, I., 2022. Brain Computer Interface for Speech Synthesis Based on Multilayer Differential Neural Networks. Cybernetics and Systems, 53(1), pp.126–140.

[38] Mantini, D., Perrucci, M. G., Del Gratta, C., Romani, G. L., Corbetta, M. (2007). Electrophysiological signatures of resting state networks in the human brain. Proceedings of the National Academy of Sciences, 104(32), 13170–13175.

[39] Mantini, D., Franciotti, R., Romani, G.L. and Pizzella, V., 2008. Improving MEG source localizations: an automated method for complete artifact removal based on independent component analysis. NeuroImage, 40(1), pp.160–173.

[40] Marino, M., Liu, Q., Samogin, J., Tecchio, F., Cottone, C., Mantini, D. and Porcaro, C., 2019. Neuronal dynamics enable the functional differentiation of resting state networks in the human brain. Human brain mapping, 40(5), pp.1445–1457.

[41] Moosmann, M., Ritter, P., Krastel, I., Brink, A., Thees, S., Blankenburg, F., Taskin, B., Obrig, H. and Villringer, A., 2003. Correlates of alpha rhythm in functional magnetic resonance imaging and near infrared spectroscopy. Neuroimage, 20(1), pp.145–158.

[42] Murphy, A.C., Bertolero, M.A., Papadopoulos, L., Lydon-Staley, D.M. and Bassett, D.S., 2020. Multimodal network dynamics underpinning working memory. Nature communications, 11(1), p.3035.

[43] Norman, K.A., Polyn, S.M., Detre, G.J. and Haxby, J.V., 2006. Beyond mind-reading: multi-voxel pattern analysis of fMRI data. Trends in cognitive sciences, 10(9), pp.424–430.

[44] Rubinov, M. and Sporns, O., 2010. Complex network measures of brain connectivity: uses and interpretations. Neuroimage, 52(3), pp.1059–1069.

[45] Samogin, J., Marino, M., Porcaro, C., Wenderoth, N., Dupont, P., Swinnen, S.P. and Mantini, D., 2020. Frequency-dependent functional connectivity in resting state networks. Human brain mapping, 41(18), pp.5187–5198.

[46] Scheeringa, R., Koopmans, P.J., van Mourik, T., Jensen, O. and Norris, D.G., 2016. The relationship between oscillatory EEG activity and the laminar-specific BOLD signal. Proceedings of the National Academy of Sciences, 113(24), pp.6761–6766.

[47] Shah, N., Arrubla, J., Rajkumar, R., Farrher, E., Mauler, J., Kops, E.R., Tellmann, L., Scheins, J., Boers, F., Dammers, J. and Sripad, P., 2017. Multimodal fingerprints of resting state networks as assessed by simultaneous trimodal MR-PET-EEG imaging. Scientific reports, 7(1), p.6452.

[48] Sherafati, A., Snyder, A.Z., Eggebrecht, A.T., Bergonzi, K.M., Burns-Yocum, T.M., Lugar, H.M., Ferradal, S.L., Robichaux-Viehoever, A., Smyser, C.D., Palanca, B.J. and Hershey, T., 2020. Global motion detection and censoring in high-density diffuse optical tomography. Human Brain Mapping, 41(14), pp.4093–4112.

[49] Shin, J., von Lühmann, A., Blankertz, B., Kim, D.W., Jeong, J., Hwang, H.J. and Müller, K.R., 2016. Open access dataset for EEG+ NIRS single-trial classification. IEEE Transactions on Neural Systems and Rehabilitation Engineering, 25(10), pp.1735–1745.

[50] Sirpal, P., Damseh, R., Peng, K., Nguyen, D.K. and Lesage, F., 2022. Multimodal autoencoder predicts fNIRS resting state from EEG signals. Neuroinformatics, 20(3), pp.537–558.

[51] Sole-Ribalta, A., De Domenico, M., Kouvaris, N.E., Diaz-Guilera, A., Gomez, S. and Arenas, A., 2013. Spectral properties of the Laplacian of multiplex networks. Physical Review E, 88(3), p.032807.

[52] Stam, C.J., Tewarie, P., Van Dellen, E., Van Straaten, E.C.W., Hillebrand, A. and Van Mieghem, P., 2014. The trees and the forest: characterization of complex brain networks with minimum spanning trees. International Journal of Psychophysiology, 92(3), pp.129–138.

[53] Strangman, G., Boas, D.A. and Sutton, J.P., 2002. Non-invasive neuroimaging using near-infrared light. Biological psychiatry, 52(7), pp.679–693.

[54] Surampudi, S.G., Naik, S., Surampudi, R.B., Jirsa, V.K., Sharma, A. and Roy, D., 2018. Multiple kernel learning model for relating structural and functional connectivity in the brain. Scientific reports, 8(1), p.3265.

[55] Tadel, F., Baillet, S., Mosher, J.C., Pantazis, D. and Leahy, R.M., 2011. Brainstorm: a user-friendly application for MEG/EEG analysis. Computational intelligence and neuroscience, 2011, pp.1–13.

[56] Tewarie, P., van Dellen, E., Hillebrand, A. and Stam, C.J., 2015. The minimum spanning tree: an unbiased method for brain network analysis. Neuroimage, 104, pp.177–188.

[57] Uludăg, K. and Roebroeck, A., 2014. General overview on the merits of multimodal neuroimaging data fusion. Neuroimage, 102, pp.3–10.

[58] Xue, W., Bowman, F.D., Pileggi, A.V. and Mayer, A.R., 2015. A multimodal approach for determining brain networks by jointly modeling functional and structural connectivity. Frontiers in computational neuroscience, 9, p.22.

[59] Van Dellen, E., de Witt Hamer, P.C., Douw, L., Klein, M., Heimans, J.J., Stam, C.J., Reijneveld, J.C. and Hillebrand, A., 2013. Connectivity in MEG resting-state networks increases after resective surgery for low-grade glioma and correlates with improved cognitive performance. Neuroimage: clinical, 2, pp.1–7.

[60] Vaiana, M. and Muldoon, S.F., 2020. Multilayer brain networks. Journal of Nonlinear Science, 30(5), pp.2147–2169.

[61] Zalesky, A., Fornito, A. and Bullmore, E.T., 2010. Network-based statistic: identifying differences in brain networks. Neuroimage, 53(4), pp.1197–1207.

[62] Zhang, F., Cheong, D., Khan, A.F., Chen, Y., Ding, L. and Yuan, H., 2021. Correcting physiological noise in whole-head functional nearinfrared spectroscopy. Journal of neuroscience methods, 360, p.109262.

[63] Zhang, Y., Wang, L., Sun, W., Green II, R.C. and Alam, M., 2011. Distributed intrusion detection system in a multi-layer network architecture of smart grids. IEEE Transactions on Smart Grid, 2(4), pp.796–808.

